# Bidirectional neuronal migration coordinates retinal morphogenesis by preventing spatial competition

**DOI:** 10.1101/2021.02.08.430189

**Authors:** Mauricio Rocha-Martins, Jenny Kretzschmar, Elisa Nerli, Martin Weigert, Jaroslav Icha, Eugene W. Myers, Caren Norden

## Abstract

While the design of industrial products is often optimized for the sequential assembly of single components, organismal development is hallmarked by the concomitant occurrence of tissue growth and organization. Often this means that proliferating and differentiating cells occur at the same time in a shared tissue environment that continuously changes. How cells adapt to architectural changes in order to prevent spatial interference remains unclear. To understand how cell movements important for growth and organization are orchestrated, we here study the emergence of photoreceptor neurons that occur during the peak of retinal growth using zebrafish, human tissue and human organoids. Quantitative imaging reveals that successful retinal morphogenesis depends on active bidirectional photoreceptor translocation. This leads to a transient transfer of the entire cell population away from the apical proliferative zone. This migration pattern is driven by distinct cytoskeletal machineries, depending on direction: microtubules are required for basal translocation, while actomyosin drives apical movement. Blocking photoreceptor translocation leads to apical overcrowding that hampers progenitor movements. Thus, photoreceptor migration is crucial to prevent competition for space and thereby allows concurrent tissue growth and lamination. This shows that neuronal migration, in addition to its canonical role in cell positioning, is involved in coordinating morphogenesis.

## Main

During organ development, the correct number of cells needs to be generated and, upon differentiation, these cells need to be spatially organized to assure tissue function. One way to achieve organogenesis would be the ‘bang-bang’ approach, a term often used in engineering to describe a system that switches abruptly between two states to minimize costs while at the same time enhancing performance ^1–3^. In development, this would entail that tissues first grow to their correct size and generate the accurate number of cells and then switch to a phase of differentiation and remodeling. However, such design strategy seem rather an exception in biological systems, for example, the intestinal crypts ^4^. In most other instances, including the formation of the pancreas, heart or brain, organogenesis involves a complex choreography during which the expansion of cell number coincides with the onset of differentiation and the establishment of functional tissue architecture ^5–7^. Thus, it is important to understand how concomitant growth and differentiation are coordinated in a shared environment.

During brain development, newborn neurons often leave their initial position and migrate to the locations at which they later function when the tissue is still proliferative and growing. Thus, migrating neurons and remaining proliferating cells need to adjust to changing tissue environment in order to prevent spatial interference. How this is achieved at the cellular and tissue level is not well understood. One part of the CNS that allows to study how tissue growth and differentiation occur synchronously is the vertebrate retina, the part of the brain responsible for visual perception. Here, the organization of retinal neurons into distinct layers ensures proper circuit formation and thereby organ function ^8,9^. Further, retinal architecture is conserved across species, allowing the comparison of findings across model systems. Recent studies have shown that in the zebrafish retina, complex migration patterns occur simultaneously at stages when the tissue undergoes significant growth ^10–13^. However, how progenitor and neuronal cell movements are coordinated in the same space has not yet been explored. Thus, we here investigated how photoreceptors (PRs) that emerge at peak proliferation stages ^10^ form the apical photoreceptor cell layer despite ongoing apical proliferation. We show that all PRs emerging apically exhibit transient basal translocation across the neuroepithelium but return apically for final positioning. This phenomenon is seen in zebrafish, human tissue and human organoids. We further demonstrate that, in zebrafish, basal and apical migration are active and driven by distinct cytoskeletal machineries: microtubules in basal direction and actomyosin in apical direction. While this bidirectional movement is dispensable for proper photoreceptor positioning, it is crucial to prevent competition for space with the apically dividing progenitors. Thereby PR translocation ensures concurrent tissue growth and lamination in zebrafish and humans, indicating that it is a fundamental aspect of retinal morphogenesis.

### Photoreceptors undergo bidirectional migration with direction-dependent kinetics

PRs are retinal neurons that are born at the same place at which they later function. However, recent studies reported a transient population of cells expressing PR markers away from the PR layer in developing zebrafish and human retinal organoids ^14–16^. To test whether this ectopic population was due to photoreceptor translocation, we imaged zebrafish retinas *in vivo* at near-physiological conditions using light-sheet microscopy^17^ and reporter genes for early neurogenic progenitors (Tg(ath5:gap-RFP)) and differentiating PRs (Tg(crx:gap-CFP)) ^18,19^. All time-lapse data was restored using the deep learning-based algorithm CARE to improve signal-to-noise ratio and allow tracing of emerging PRs (Extended Data Fig. 1A; for details, see Methods) ^20^. Imaging of whole retinas revealed that, upon apical birth, both daughter cells of Ath5+ neurogenic divisions moved basalward across the neuroepithelium (Fig. 1A, Supplementary Video 1). At basal positions, one of the sister cells that exhibited elongated unipolar morphology (Fig. 1B) started to express crx:CFP and thereby identified as an emerging cone PR (Fig. 1A, Supplementary Video 1). As development progressed, all Crx positive (from here on Crx+) cells returned to the apical surface (Fig. 1A, C and C’, Supplementary Video 2).

**Fig. 1.**
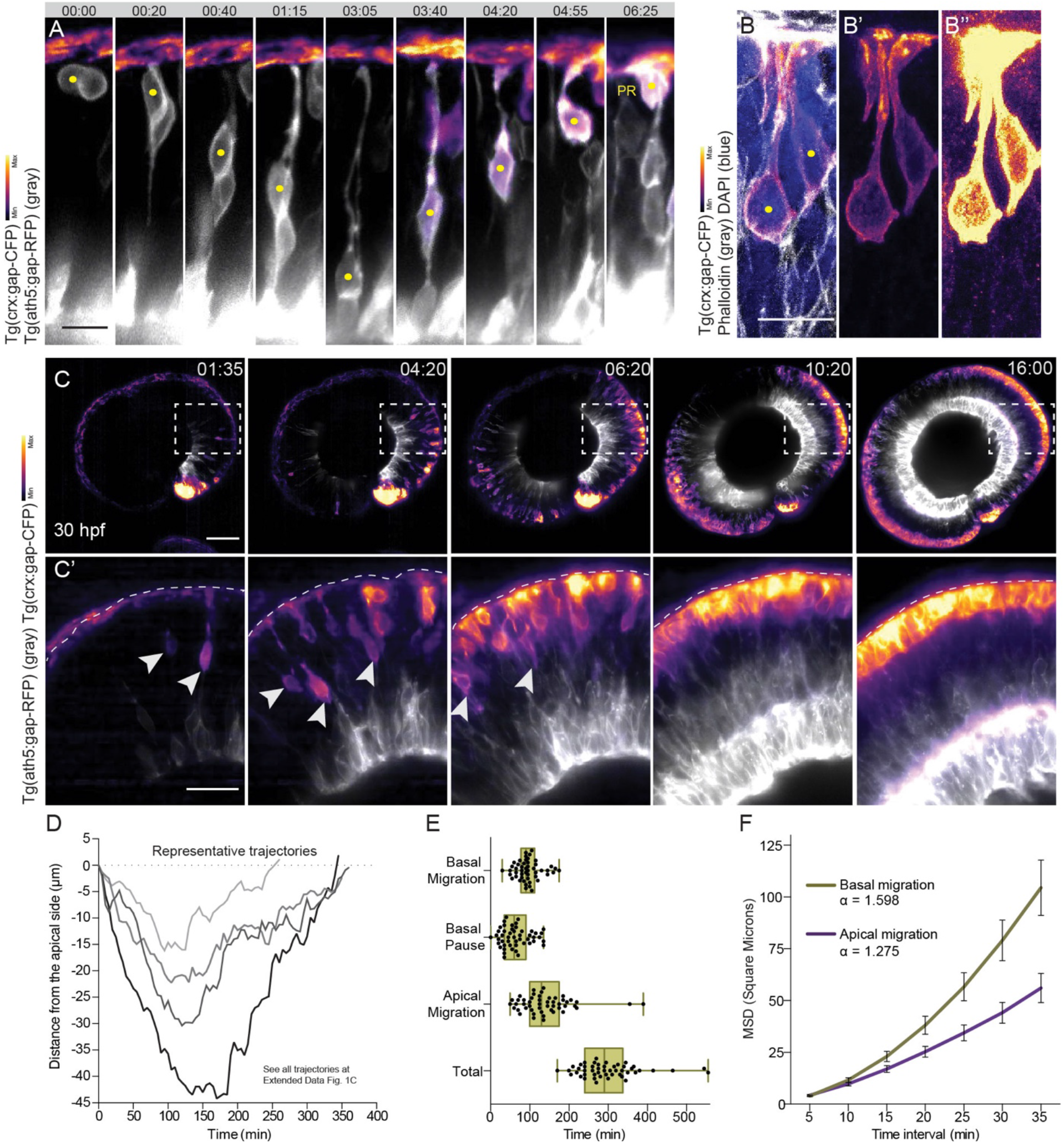
Emerging photoreceptors undergo bidirectional translocation in the zebrafish retina with direction-dependent kinetics. (A) Bidirectional migration of photoreceptors (PR). Tg(ath5:gap-RFP) (gray) labels early neurogenic progenitors; Tg(crx:gap-CFP) labels PRs. Yellow dots mark PRs. (B) PR morphology at 42 hpf. Staining: Tg(crx:gap-CFP) labels PR. Phalloidin (gray) labels F-actin; DAPI (blue) labels nuclei. B’ and B’’ show crx:CFP signal with different adjustments of brightness and contrast to highlight the absence of a basal attachment. Yellow dots mark PRs. (C) Time-series of PR lamination in the full retina. Tg(ath5:gap-RFP) (gray) labels early neurogenic progenitors; Tg(crx:gap-CFP) labels PRs. Dashed white boxes indicate areas shown in C’. Arrowheads mark migrating PRs. (D) Representative trajectories of PR bidirectional migration from birth to final positioning relative to the apical surface (0 μm). (E) Duration of the different phases of PR migration shown as boxplot with superimposed individual measurements (dots; n= 49 cells; N= 17 embryos). The lower and upper hinges of the box indicate the 25th and 75th percentile, respectively; middle line indicates the median, whiskers indicate min and max. P-value for comparison between apical and basal migration using Wilcoxon matched-pairs signed rank test: < 0.0001. (F) Mean square displacement of PR’s basal and apical migration shown as mean of all tracks ± SEM (n= 33 cells; N= 15 embryos). Scale bars: 10 μm (A-B), 20 μm (C’), 50 μm (C). (A-C) Lookup table shows min and max signal values. (A, C) Time is displayed as hours:minutes.

To backtrack PRs (Fig. 1D and Extended Data Fig. 1C) and quantify their movements, cells were mosaically labelled using an Ath5-driven reporter construct (ath5:GFP-CAAX; Extended Data Fig. 1B) and PRs were identified based on morphology and final position ^21^. All emerging PRs exhibited a similar pattern of bidirectional translocation (n=49 cells, N=17 embryos) (Fig. 1D and Extended Data Fig. 1C). This migration pattern was also seen for L-cone photoreceptors labeled by a trβ2 promoter (trβ2:tdTomato; Extended Data Fig. 1E, Supplementary Video 1) ^22^.

Separating apical and basal movement phases showed that basal movement was more efficient than apical movement (Fig. 1D; Supplementary Video 1) and that apical movement was more variable in duration (Fig. 1E). Further, basally moving PRs displayed more continuous and monotonic movements while apically moving PRs translocated in a more saltatory manner (Fig. 1D). Directionality ratio and mean square displacement (MSD) analysis confirmed these notions (Fig. 1F and Extended Data Fig. 1D). Nevertheless, both movements occurred in a directed manner, as shown by the positive curvature of the MSD analysis (Fig. 1F).

Thus, emerging PRs undergo bidirectional migration prior to final apical positioning and basal and apical movements display different kinetics.

### Photoreceptor migration is conserved in humans

As retinal architecture is highly conserved in many vertebrates ranging from fish to mammals, we asked whether the bidirectional migration of emerging PRs seen in zebrafish is a fundamental aspect of retinal development by studying PR behavior in a distantly related species, humans. To this end, we stained human fetal retinas (provided by the human tissue bank (HDBR)) against Otx2 and Crx, two transcription factors observed during PR fate determination and differentiation, respectively ^23,24^(Fig. 2A).

**Fig. 2.**
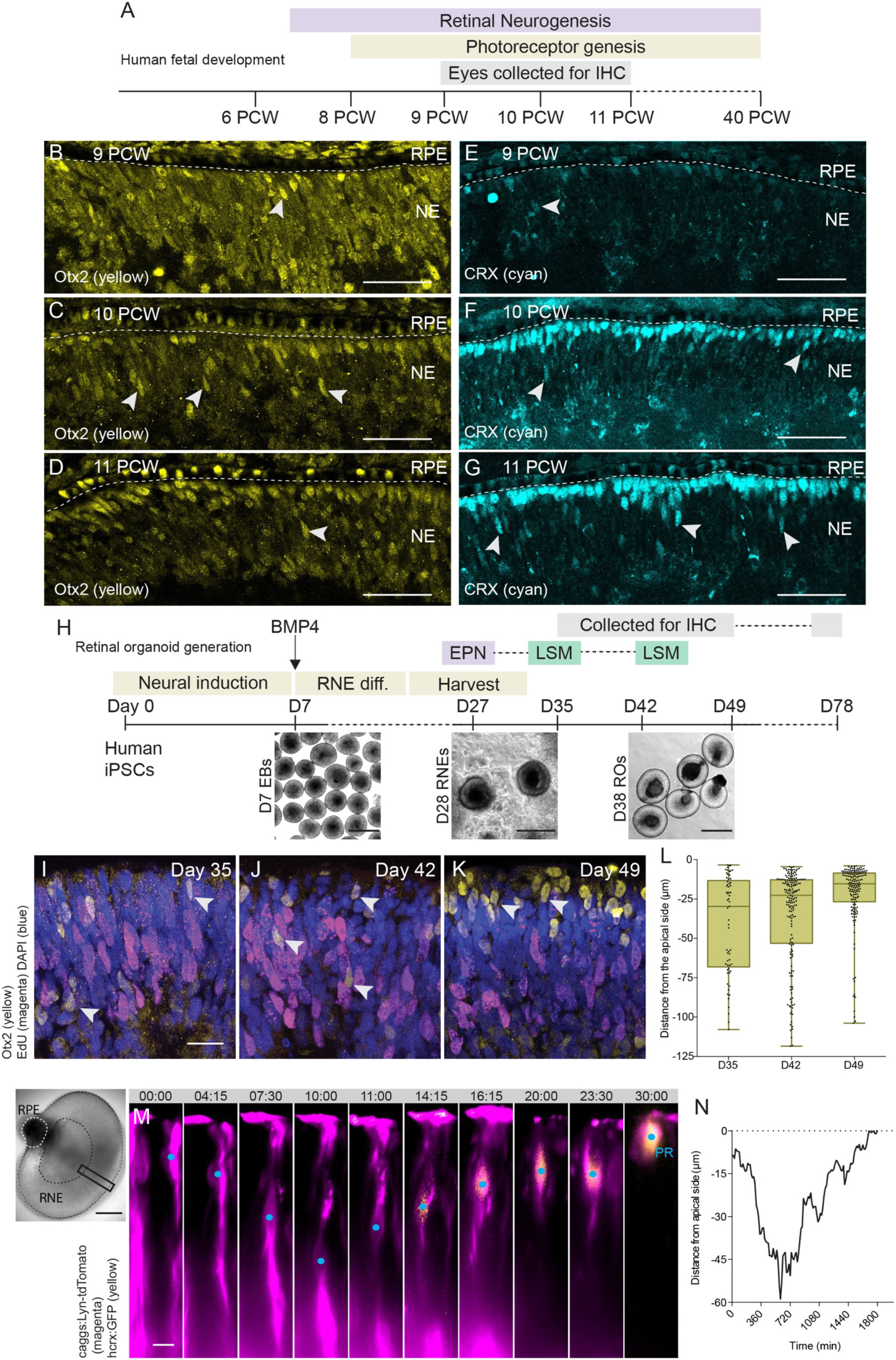
Bidirectional photoreceptor translocation is conserved in human tissue and human organoids. (A) Timeline of analysis of PR positioning in human fetal eyes. (B-G) Distribution of PRs along the apical-basal axis of the neuroepithelium of developing human fetal retinas. Staining: Otx2 (yellow) labels Otx2+ PRs; CRX (cyan) labels CRX+ PRs. (H) Schematic of human retinal organoid protocol (see Material and Methods). (I-K) Spatial distribution of Otx2+ PRs along the apical-basal axis of the neuroepithelium of developing human retinal organoids. Staining: Otx2 (yellow) labels Otx2+ PRs; EdU (magenta) labels cells in S-phase; DAPI (blue) labels nuclei. Arrowheads mark Otx2+ PRs. (L) Boxplot of distance of Otx2+ cells from the apical surface. The lower and upper hinges of the box indicate the 25th and 75th percentile, respectively; middle line indicates the median, whiskers indicate min and max. Dots represent individual measurements. D35: n= 61 cells, N= 4 organoids; D42: n= 169 cells, N= 3 organoids; D49: n= 157 cells, N= 4 organoids. P-value for Dunn’s Kruskal-Wallis Multiple Comparisons with Holm-Bonferroni correction: 0.403282 for D35 and D42; 0.000092 for D35 and D49; 0.000030 for D42 and D49. (M) Retinal organoid at day 33 of culture mounted for LSM imaging. Scale bar: 200 μm. Black and white dashed lines delineate the retinal neuroepithelium and the retinal pigment epithelium, respectively. Black box indicates the approximate region displayed in the montage of photoreceptor translocation. hcrx:GFP (yellow) labels PR; caggs:Lyn-tdTomato (magenta) labels transfected cells. Time is displayed in hours:minutes. Blue dots mark PRs. (N) Trajectory of a representative PR positive for hcrx-driven GFP relative to the apical surface (0 μm). Scale bars: 10 μm (M), 20 μm (I-K), 50 μm (B-G), 500 μm (H).

At 9 weeks post-conception (PCW), only few Crx+/Otx2+ cells were observed at the apical surface of the central retina while other cells occupied positions along the apicobasal axis (Fig. 2B and E). At 10 PCW and 11 PCW, however, an increasing number of Crx+/Otx2+ PRs was detected along the entire thickness of the neuroepithelium (Fig. 2C-G). As the human retina differentiates in a central-to-peripheral gradient, different areas of the 11 PCW retina were analysed as a proxy for developmental progression. Otx2+ cells at the periphery occupy both apical and basal positions, while in central, more developed regions, cells became more restricted to the apical surface (Extended Data Fig. 2A). Thus, the spatial distribution of emerging PRs during human retinogenesis was similar to what was seen in zebrafish.

To be able to follow these findings in more detail, we turned to human retinal organoids derived from human iPSCs using a modified version of previously established protocols (Fig. 2H) ^25,26^. Organoids were analysed for the spatial distribution of emerging PRs from the onset of differentiation at day 35 until early stages of lamination at day 78 (Fig. 2H). Postmitotic photoreceptors were identified based on the expression of Otx2 and the absence of EdU signal (all Otx2+ cells (n=387) were negative for EdU). At early stages of organoid differentiation (day 35 and day 42), Otx2+ cells were found along the entire thickness of the neuroepithelium (Fig. 2I-J, L), similar to what was seen in zebrafish and human tissue. At Day 49 these cells became increasingly restricted to the apical surface (Fig. 2K-L). At early lamination stages (Day 78), PRs were found at apical positions as shown by recoverin (PR marker) and HuC/D (basal neuron marker) staining (Extended Data Fig. 2B).

To test whether this spatial distribution is due to bidirectional PR translocation as in zebrafish, a protocol for live imaging of intact organoids using light-sheet microscopy was established (see Methods). In brief, retinal organoids were electroporated prior to differentiation (day 27) with a transfection reporter (caggs:Lyn-tdTomato) and a human Crx-driven reporter construct (hcrx:GFP) to label the emerging PRs (Fig. 2H). The majority of the GFP+ cells expressed endogenous Otx2 at day 43 (Extended Data Fig. 2C), confirming the specificity of the reporter. At days 33 and 43 (6 and 16 days after electroporation), organoids were imaged live every 15 min for 18-35 hours (Fig. 2H, Supplementary Video 3). Also in human organoids, emerging PRs undergo bidirectional translocation (Fig. 2M-N, Extended Data Fig. 2D and Supplementary Video 3; n= 7 cells, N= 2 organoids). However, due to different developmental timing and thickness of the organoid neuroepithelium, the human organoid PRs translocated over longer distances and longer time periods (Fig. 2N and Extended Data Fig. 2D). Nevertheless, the overall trajectories were strikingly similar to zebrafish, as in both systems basal movement was continuous, whereas the apical translocation was saltatory (Fig. 2N and Extended Data Fig. 2D).

These results show that PR translocation is comparable in retinas of systems as dissimilar as zebrafish and humans indicating that it is an important feature for retinal development.

### Stable microtubules are required for active basal photoreceptor migration but dispensable for apical migration

We noted kinetic differences between basal and apical movements of emerging PRs in both zebrafish embryos and human organoids. Thus, we turned back to the zebrafish system to better understand the cellular mechanisms that drive the PR translocation in the different directions. The kinetics of initial basal translocation of the emerging PRs were strikingly similar to what has been previously reported for their sister cells, RGCs ^11^ (Fig. 3A and Extended Data Fig. 3A). Despite the close proximity of the sister cells, RGCs are dispensable for basal translocation of emerging PRs, as they still translocate when RGCs are genetically depleted using an established Ath5 morpholino ^27^ (Fig. 3B and Extended Data Fig. 3B). Furthermore, some emerging PRs in control embryos were able to move to basal positions even when a considerable distance between sister cells was observed (Fig. 3C). These findings indicated that PRs moved autonomously with very similar kinetics to sister RGCs. As basal RGC translocation depends on a stabilized microtubule cytoskeleton ^11^, we tested whether a similar mechanism was at play for emerging PRs. To this end, the distribution of the microtubule-associated protein Doublecortin (βactin:GFP-DCX) was analysed during basalward PR movements. Microtubules were enriched in the apical process shortly after their birth and during basal movement (Fig. 3D). Acetylated tubulin staining showed that these apical microtubules were stabilized as previously seen in RGCs (Fig. 3E) ^11^.

**Fig. 3.**
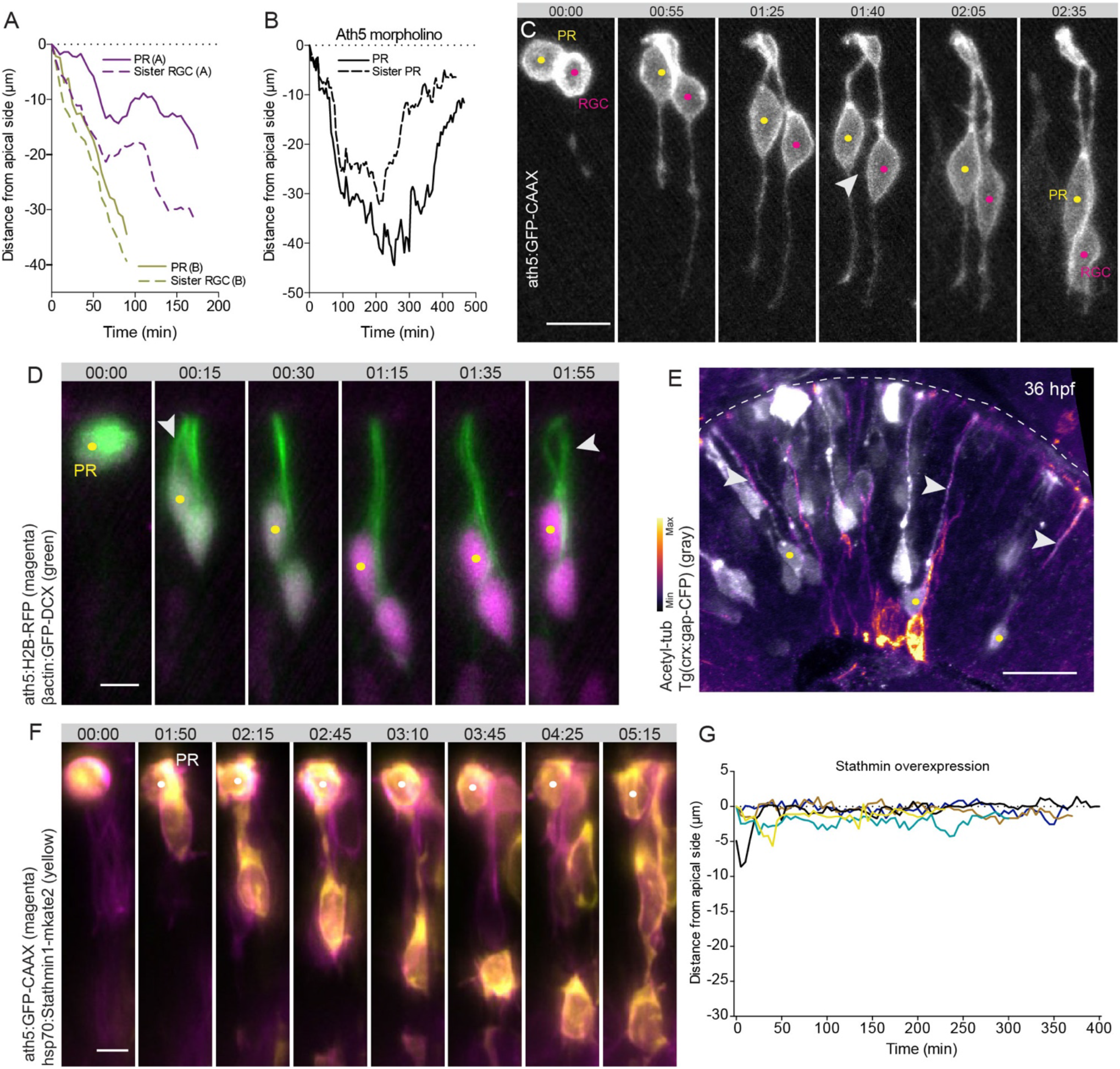
Stable microtubules are required for basal PR migration but dispensable for apical migration. (A) Trajectories of basal migration of two representative PRs and the respective sister RGCs relative to the apical surface (0 μm). (B) Trajectories of bidirectional migration of representative sister PRs relative to the apical surface (0 μm) upon Ath5 morpholino-mediated knockdown. (C) Basal migration of PR and sister RGC showing that sister cells are not in direct vicinity. Yellow dots mark PR; magenta dots mark RGC; arrowhead marks space between sister cells. (D) Microtubule (MT) distribution during PR basal migration. ath5:H2B-RFP (magenta) labels early neurogenic progenitors; βactin:GFP-DCX (green) labels MTs. Yellow dots mark PR; arrowheads mark MTs in the apical process. (E) Distribution of stable MTs in basally migrating PRs at 36 hpf. Staining: Tg(crx:gap-CFP) (gray) labels PRs; Acetylated tubulin labels stable MTs, lookup table shows min and max signal values. Yellow dots mark PRs; arrowheads mark apical process of PRs. (F-G) Effect of Stathmin1 overexpression on PR migration. (F) ath5:GFP-CAAX (magenta) labels early neurogenic progenitors; hsp70:Stathmin1-mKate2 (yellow). White dots mark PR. (G) Trajectories of PRs upon Stathmin1 overexpression relative to the apical surface (0 μm). Scale bars: 5 μm (D, F), 10 μm (C), 20 μm (E). (C-D, F) Time is displayed in hours:minutes.

To test the active involvement of these stabilized microtubules in basal PR translocation, microtubules were depolymerized by colcemid. This resulted in a premature accumulation of emerging PRs at the apical surface (Extended Data Fig. 3D). This finding was substantiated by overexpression of the microtubule-destabilizing protein Stathmin1 ^11^ which lead to erratic movements of emerging PRs and their premature return to the apical surface (Extended Data Fig. 3E-F, Supplementary Video 4). In severe cases, a complete stalling of PRs at the apical surface was observed (Fig. 3F-G, Supplementary Video 4).

Interestingly, colcemid induced microtubule depolymerization did not similarly affect apical movement of emerging PRs (Extended Data Fig. 3G-H, Supplementary Video 5) and no enrichment of acetylated tubulin within the apical process was seen when cells returned apically (42 hpf) (Extended Data Fig. 3C). Together, these results argue that basal translocation of emerging PRs is microtubule-dependent, while their return to the apical surface is not.

### Actomyosin is the main driver of photoreceptor apical migration

In addition to microtubules, the actin cytoskeleton has been shown to be responsible for cell and organelle translocation in neuroepithelial and neuronal cells ^28–30^. Thus, we explored whether actomyosin could be involved in apical translocation of emerging PRs. To this end, crx:gap-CFP expressing retinas were stained for F-actin and active myosin at 42 hpf, when most emerging PRs undergo apical translocation. In some CRX+ cells, enrichment of F-actin and active myosin was observed basal to the nucleus (Fig. 4A and B). Live imaging of Utrophin (ath5:GFP-Utrophin) revealed that actin distribution was dynamic, showing no difference between basal and apical translocation (Extended Data Fig. 4A). In contrast, Myosin distribution (Ath5:MRLC2-mKate2) that colocalized with actin (Fig. 4F), changed depending on translocation directionality and was seen to be basally enriched specifically when emerging PRs returned to the apical surface (Fig. 4C, Supplementary Video 6, n=18 cells/ 6 embryos). While these myosin enrichments were transitory (Fig. 4D-D’), quantification confirmed that they were reproducibly more prevalent basally (Fig. 4E, n= 8 cells/ 4 embryos).

**Fig. 4.**
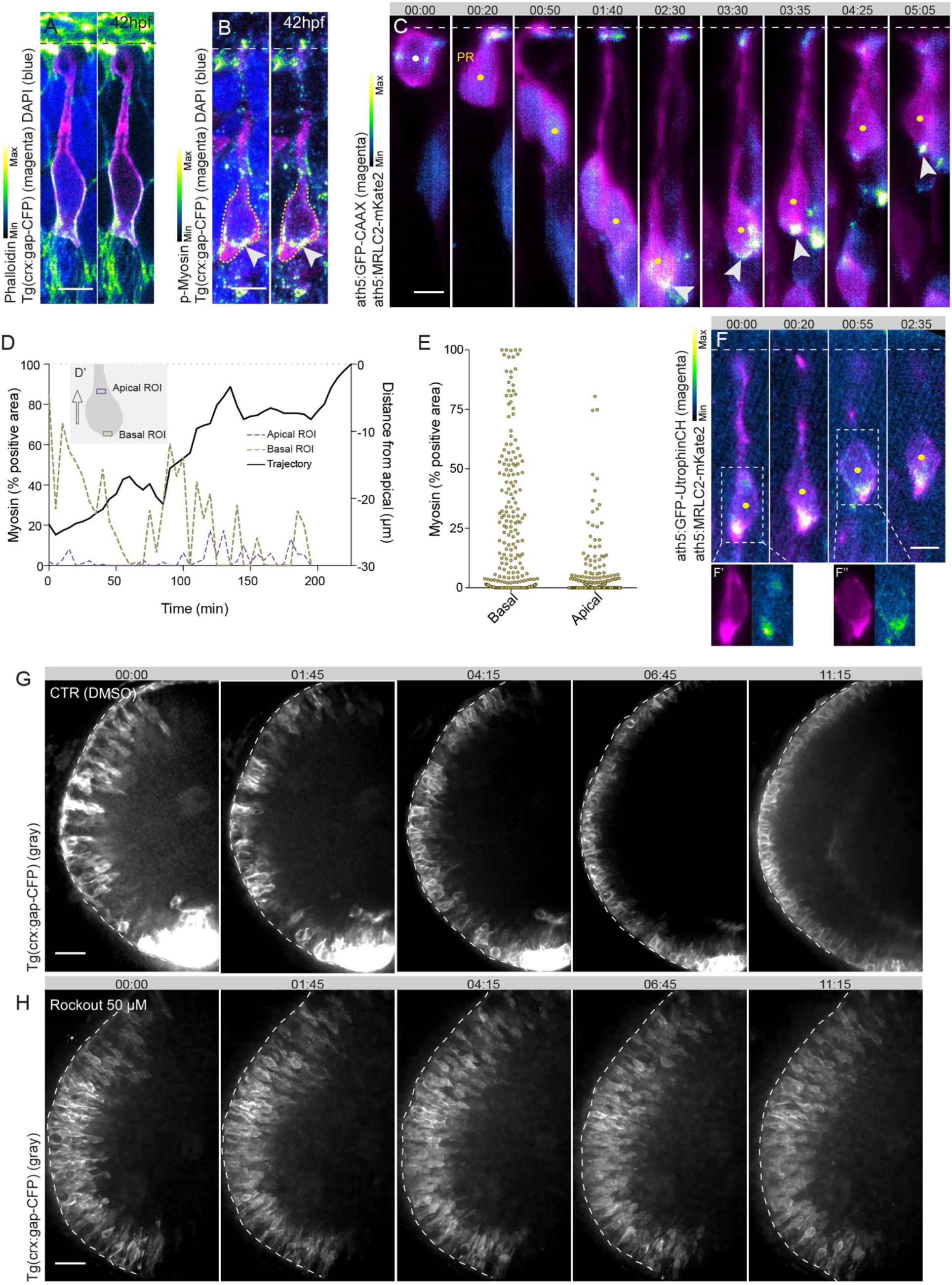
Actomyosin is the main driver of apical migration of photoreceptors. (A) F-actin distribution in migrating photoreceptor (PR) at 42 hpf. Staining: Phalloidin labels F-actin; Tg(crx:gap-CFP) (magenta) labels PR; DAPI (blue) labels nuclei. (B) Active myosin distribution in migrating PR at 42 hpf. Staining: p-myosin labels active myosin; Tg(crx:gap-CFP) (magenta) labels PR; DAPI (blue) labels nuclei. Dashed yellow line outlines PR; arrowhead marks myosin enrichment. (C) Myosin distribution during PR migration. ath5:GFP-CAAX (magenta) labels early neurogenic progenitors; ath5:MRLC2-mKate2 labels myosin. White dot marks apical division; yellow dots mark PR; arrowheads mark myosin enrichment. (D) Myosin enrichments basal and apical to the nucleus over time during apical migration of a representative PR (ROIs depicted in D’). (E) Scatter plot quantification of myosin enrichments basal and apical to the nucleus during apical migration of PRs (n= 8 cells, N= 4 embryos, 218 timepoints). P-value for Wilcoxon matched-pairs signed rank test: < 0.0001. (F) Myosin and actin distribution during apical PR translocation. ath5:Utrophin-GFP (magenta) labels actin in early neurogenic progenitors; ath5:MRLC2-mKate2 labels myosin early neurogenic progenitors. Yellow dots mark PR; Dashed white boxes outline areas shown in F’ and F’’. (G-H) Time-series of retinas treated with the actomyosin activity inhibitor Rockout (50μM, H) or DMSO (control, G). Tg(crx:gap-CFP) (gray) labels PRs. Scale bars: 5 μm (A-C,F), 20 μm (G-H). (A-C, F) lookup table shows min and max signal values. (C, F-H) Time is displayed in hours:minutes.

To confirm that actomyosin contractility drives apical movement of emerging PRs, myosin was inhibited by Blebbistatin which fully abolished their apical movements (Extended Data Fig. 4C-D, Supplementary Video 5). We further tested which of the serine/threonine kinases, MLCK or ROCK, acted upstream of Myosin II activation. Transcriptome analysis showed that only low levels of MLCK were expressed in emerging PRs, while the expression of Rock genes was more prominent (Extended Data Fig. 4B). Accordingly, interference with Rock activity using Rockout (Fig. 4G-H, Supplementary Video 5) resulted in impaired apical movements, similar to inhibition by Blebbistatin (Extended Data Fig. 4C-D).

Thus, apical migration of emerging PRs is regulated by Rho-ROCK-dependent myosin contractility. Moreover, the two phases of PR migration not only exhibit different kinetics (Fig.1E-F) but are also driven by different cytoskeletal elements.

### Photoreceptor migration prevents overcrowding of the apical mitotic zone

The fact that bidirectional PR movement is conserved in zebrafish and humans and is highly regulated at the molecular level argued that, despite being counterintuitive, it played an important role for tissue formation.

Reassessment of the trajectories gathered in Fig. 1 revealed that emerging PRs stay away from the apical surface for an average of 300.6 minutes (coefficient of variation (CoV), 25.97%) while the depth of migration varied between 8.7 μm and 44.2 μm (CoV, 31.14%) (Extended Data Fig. 5A). However, depth of migration and duration away from the apical surface did not strongly correlate (Extended Data Fig. 5A) arguing that moving PRs out of the way for a certain amount of time was more relevant than moving them as far basally as possible. Furthermore, PR migration did not seem to be essential for the determination or maintenance of cellular fate as when migration away from the apical surface was perturbed by Stathmin overexpression, emerging PRs nevertheless positioned correctly and showed typical PR morphology (Fig. 3F). Thus, other non-cell autonomous prerequisites were more likely responsible for the migration patterns observed.

**Fig. 5.**
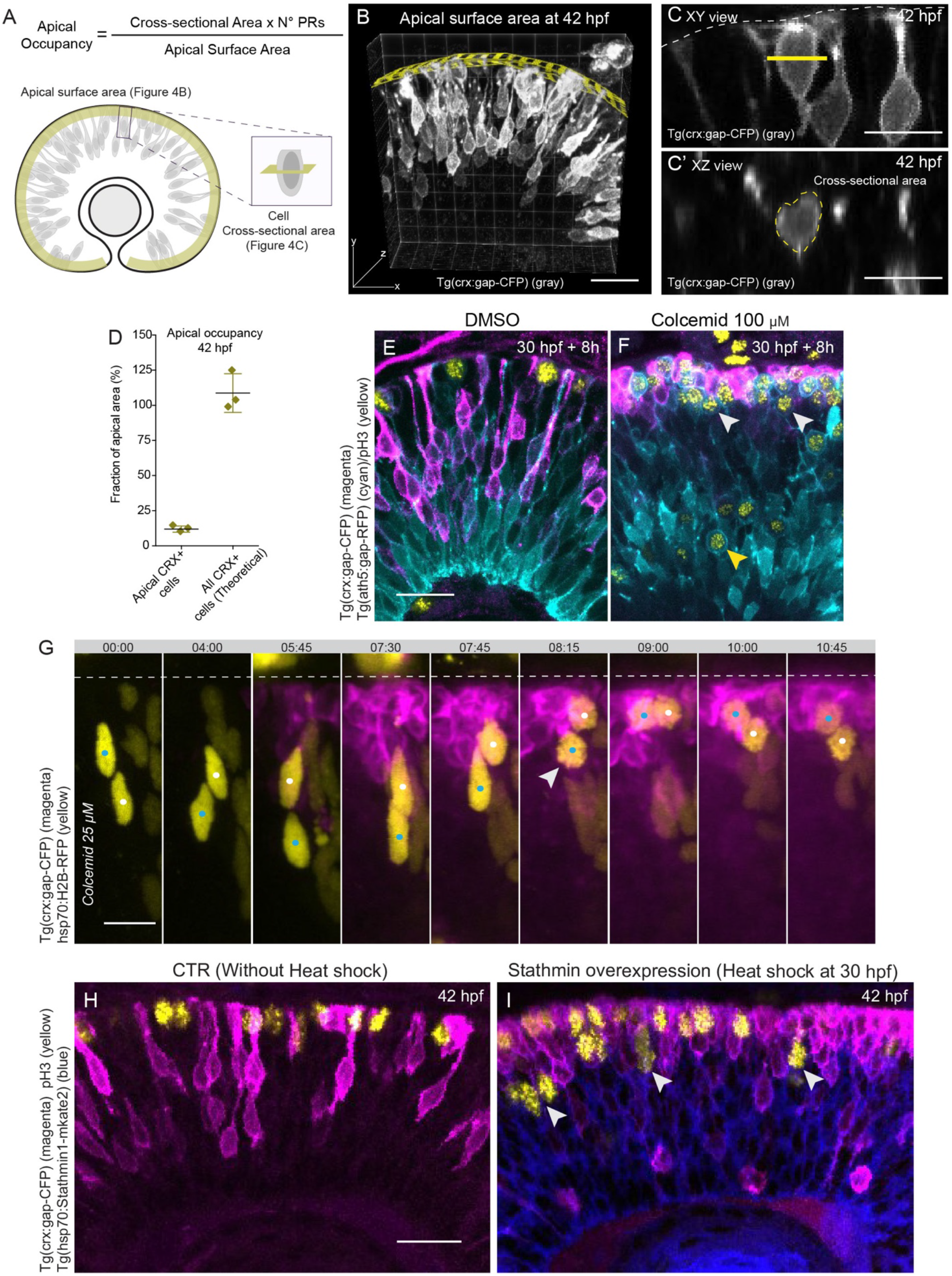
Photoreceptor migration prevents overcrowding of the apical mitotic zone. (A) Scheme of apical occupancy analysis. (B) Measurement of apical surface area of retinal tissue at 42 hpf using FIJI plugin Volume Manager. Tg(crx:gap-CFP) (gray) labels photoreceptors (PRs). Yellow grid marks the apical surface. (C) Measurement of the cross-sectional area of apical PRs at 42 hpf. Tg(crx:gap-CFP) (gray) labels photoreceptors (PRs). Yellow line in the XY view outlines the plane shown in XZ view (C’). (D) Fraction of apical area occupied by CRX+ cells (measured) versus all CRX+ cells (theoretical) at 42hpf shown as mean ± SD (N=3 embryos). (E-F) Effect of colcemid-induced blockage of PR migration on progenitor divisions. Embryos were treated with 100 μM of colcemid (F) or DMSO (control E) from 30 hpf for 8h. Staining: Tg(crx:gap-CFP) (magenta) labels PRs; Tg(ath5:gap-RFP) (cyan) labels RGCs, PRs and neurogenic progenitors; pH3 (yellow) labels mitotic cells. White arrowheads mark subapical mitotic cells; yellow arrowhead marks basal mitotic cell. (G) Subapical mitoses upon colcemid-induced blockage of PR migration. Embryos were treated with 25 μM colcemid. Tg(crx:gap-CFP) (magenta) labels PRs; hsp70:H2B-RFP (yellow) labels apically migrating nuclei. Time is displayed in hours:minutes. Blue and white dots mark progenitor cells; arrowhead marks subapical mitosis. (H-I) Effect of Stathmin-induced blockage of PR migration on progenitor divisions at 42 hpf. Heat shock induction of Stathmin overexpression at 30 hpf. Staining: Tg(crx:gap-CFP) (magenta) labels PRs; pH3 (yellow) labels mitotic cells; Tg(hsp70:Stathmin1-mkate2) (blue) labels Stathmin. Arrowhead marks ectopic mitosis. Scale bars: 10 μm (C, G), 20 μm (B, E, F, H, I).

As PRs are one of the first neurons to be born in the developing retina ^31^, their generation coincides with a last peak of proliferative growth ^10^. Indeed, a substantial number of actively cycling progenitors is still present when PRs start to emerge as confirmed by EdU labelling in zebrafish (Extended Data Fig. 5B) as well as in human organoids (Fig. 2I-K).

During this proliferative peak, all progenitor divisions occur apically and non-apical divisions disturb tissue integrity ^32^. Taking these notions into consideration, we asked whether basal translocation of emerging PRs could be linked to the need for apical surface availability during the proliferative peak. We analysed the fraction of the apical surface that was occupied by apical photoreceptors at this stage (Fig. 5A). To this end, we quantified tissue surface area (Fig. 5B), the average cross-sectional area of apical PRs (Fig. 5C) and the total number of emerging PRs (Extended Data Fig. 5C). This showed that only 11,2% of PRs were located at apical positions at 42 hpf occupying 11,9% ± 2,2% (mean ± SD) of the apical surface area (Fig. 5D). In contrast, in a theoretical scenario in which all PRs born at 42 hpf would not undergo translocation, the apical occupancy would rise to 108,7% ± 13,8% (mean ± SD) (Fig. 5D). We thus speculated that overcrowding of the apical surface could potentially affect incoming progenitor divisions if emerging PRs were not moving out of the way. We tested this notion by inducing a tissue-wide arrest of basal translocation of emerging PRs using colcemid (Extended Data Fig. 3D). This led to PRs occupying the complete apical surface of the tissue (Fig. 5E-F). Consequently, progenitor divisions took place at subapical and basal positions (Fig. 5E-F). This finding was substantiated by live imaging experiments of colcemid treated embryos which revealed that PRs indeed prematurely congested the apical surface (Fig. 5G). This resulted in progenitor cells that attempted to move their nuclei apically but encountered a physical barrier in form of the prematurely formed PR layer. In turn, progenitor cells entered mitosis at subapical positions (Fig. 5G, Supplementary Video 7). In regions without an already fully occupied PR layer, progenitor cells entered mitosis apically (Extended Data Fig. 5D, Supplementary Video 7) showing that subapical mitosis was not caused by microtubule destabilization per se, but indeed by PRs congesting the apical surface.

To further substantiate these findings, we generated a stable transgenic line harboring a heat shock-inducible stathmin (Tg(hsp70:Stathmin1-mkate2). This line reproduced the PR congestion phenotype (Extended Data Fig. 5E-F, Supplementary Video 5) and led to subapically dividing progenitor cells (Fig. 5H-I). Consistent with previous findings that apical crowding can induce progenitor delamination ^32^, blocking emerging PR translocation sometimes resulted in neuronal lamination defects and mitotic cells seen within the basal neuronal layers (Extended Data Fig. 5G-H)

Thus, the transient basal translocation of emerging PRs aids to prevent the overcrowding of the apical surface. It thereby allows progenitor cells to undergo mitosis at apical positions even at stages when differentiation and lamination are already under way.

## Discussion

This study shows that active bidirectional migration of emerging PRs in zebrafish and humans is important to coordinate growth and lamination during retinogenesis. We reveal that movements are driven by different molecular motors depending on direction: microtubules in basal direction and actomyosin in apical direction. More precisely, we propose that microtubule polymerisation propels the soma of emerging PRs in the basal direction while concurrent stabilization of microtubules prevents backward movements (Extended Data Fig. 6). In the apical direction, actomyosin contractions regulated by the Rho-ROCK pathway drive the saltatory movement of emerging PRs (Extended Data Fig. 6). While the interplay between actomyosin and microtubules to integrate the movement of different parts of a cell and the different steps of migration has already been suggested in other contexts ^33,34^, the use of two distinct cytoskeletal elements by the same emerging neuron to move in different directions has to our knowledge not yet been revealed. Next, we need to unveil the molecular and tissue wide triggers that induce the reversal of migration, an exciting topic for future investigations.

The fact that we discovered a new migratory mode for one the most widely studied neuronal cell types, the PRs ^23,35,36^, demonstrates that neuronal migration is far from being understood. Thus, studying different areas of the central nervous system will be essential to refine our understanding of the diversity of neuronal migration phenomena and how they contribute to the generation of the unique cytoarchitectures of the brain.

Generally, neuronal migration has so far mainly been recognized as an important and widespread phenomenon that ensures correct neuronal positioning. This in turn safeguards the formation of functional neural circuits. Not surprisingly, neuronal migration defects can lead to a variety of neurological disorders ^37,38^. However, our study shows that even neurons that are born at the same position at which they later reside and function, like the retinal PRs, can undergo translocation before final positioning. While at first sight a translocation resulting in a zero-net displacement seems redundant, it turns out to be essential for overall successful retinal development. Unlike other neuronal translocation phenomena ^11,13,39^, this migration is not a prerequisite for correct positioning of the cells. Instead, it is important to prevent local crowding which can induce mechanical stress that in turn can lead to progenitor delamination as seen also at earlier developmental stages ^40,41^. Thus, we propose that by preventing the competition for space, migration of the emerging PRs is crucial for the spatiotemporal coordination of tissue growth and lamination. We speculate that when neuronal tissues start to differentiate and remodel while still under active proliferation, strategies need to be deployed that nevertheless ensure apical surface availability for progenitors. In the retina, this takes the form of basal translocation of apically emerging neurons. Thus, neuronal migration in addition to its canonical role for correct cell positioning can be an important mechanism to coordinate cell and tissue wide phenomena during brain morphogenesis. As simultaneous tissue growth and differentiation are a hallmark of many developing organs, it is tempting to speculate that similar relocation strategies of entire cell populations could also be at play elsewhere.

Our finding that this phenomenon is conserved in zebrafish, human tissue and human-derived organoids suggests that it is fundamental to retinal development across evolution. It also argues that retinal morphogenesis, like tissue architecture and gene regulatory networks, is highly conserved. This area of conservation is however much less understood and more comparative studies are needed to probe how tissue growth and differentiation are orchestrated in the developing embryo and how defects of such coordination can lead to a diseased state.

## Material and Methods

### Human primary tissue

Human fetal eyes at 9 PCW (n=1), 10 PCW (n=2) and 11 PCW (n=1) were provided fixed in 4% paraformaldehyde (PFA) by the Joint MRC/Wellcome Trust (grant# 099175/Z/12/Z) Human Developmental Biology Resource (http://hdbr.org).

### Human iPSC culture and generation of retinal organoids

Human iPSCs (IMR90 clone 4 from WiCell) ^42^ were cultured with mTeSR1 medium on Matrigel (Corning) coated 6-well plates (Nunclon delta, Thermo Fisher Scientific). Cells were passaged at approximately 80% confluence using ReleSR (Stem Cell) dissociation reagent and stocks were cryopreserved in mFreSR (Stem Cell). To generate retinal organoids, we adapted previously published protocols ^25,26^. On day 0 (D0), iPSCs colonies were lifted mechanically with a cell scraper (Thermo Fisher Scientific) after Dispase (Stem Cell) treatment. The cell aggregates were further dissociated into smaller clusters by gentle pipetting through 1 mL pipette tips. Embryoid body (EB) formation was induced by culturing cell aggregates in suspension from 1-well of a 6-well plate on a 35×10mm petri dish (Thermo Fisher Scientific) with mTeSR1 medium. The aggregates were gradually transitioned into Neural-Induction Medium (NIM: DMEM/F12 1:1 supplemented with 1x GlutaMax, 1× N-2, 1x MEM Non-essential Amino Acid Solution (NEAA) and 2 μg/ml of heparin). Over the course of the first 4 days in culture, the medium was replaced with different ratios of mTeSR1/NIM: 3:1 on D1, 1:1 on D2 and 100% NIM on D3. Aggregates were transferred daily to 15 mL conical tubes with 1 mL pipette tips for medium exchange until D5, promoting dissociation of bigger aggregates and ensuring homogeneous size of the EBs. On D6, 200 EBs were selected and fresh NIM supplemented with 1.5 nM BMP4 (R&D Systems) was added to promote specification into neural retina fate ^25,43^. On D7, the selected EBs were transferred to a 35×10mm petri dish coated with Matrigel. To prolong the effects of BMP4, medium exchange was performed only on D9, D12 and D15, wherein half of the medium was replaced with NIM. On D16, the medium was fully replaced with Retinal Differentiation Medium (RDM: DMEM/F12 3:1 supplemented with 2% B-27 without vitamin A, 1x GlutaMax, 1x NEAA, 1% Penicillin-Streptomycin) and exchange was performed daily until D25-30 when the optic vesicles became visible. One day before dissection of the optic vesicles, a polytetrafluoroethylene mold made by milling was used to cast 36 hemispherical wells on a thin layer of 2% agarose (Cell culture grade, Sigma-Aldrich) in ddH_2_O in a 35×10mm petri dish. After solidification of the agarose, the dish was filled with Retinal Organoid Medium (ROM: DMEM/F12 3:1 supplemented with 2% B-27, 5% FBS, 1x GlutaMax, 1x NEAA, 0.1% Lipid concentrate, 1% Penicillin-Streptomycin and 100 μM Taurine) and maintained in the cell culture incubator until organoid dissection. The optic vesicles were dissected under a widefield microscope using 5x/0.12 or 10x/0.25 NPLAN objectives (LEICA), and 27-gauge needles (BD). Organoids were transferred to the agarose bed with fresh ROM medium and placed in individual wells to prevent fusion. From this point on, the ROM medium was exchanged every third day. In all stages of the protocol, iPSCs and developing organoids were cultured at 37°C and 5% CO_2_ in a humidified incubator (Thermo Fisher Scientific).

### Zebrafish husbandry

Wild-type zebrafish and transgenic lines were maintained at 26°C. Embryos were raised at 28.5 or 32°C in E3 medium. Medium was changed daily and supplemented with 0.2 mM 1-phenyl-2-thiourea (Sigma-Aldrich) from 8 hours post fertilization (hpf) to prevent pigmentation. Staging was performed in hours post fertilization ^44^. All animal work was performed in accordance with European Union directive 2010/63/EU, as well as the German Animal Welfare Act.

### Zebrafish transgenesis and transgenic lines used

To visualize emerging photoreceptors, Tg(ath5:gap-RFP), Tg(ath5:gap-GFP) and Tg(crx:gap-CFP) lines were used ^18,19^. To generate a stable transgenic line containing heat-shock inducible human Stathmin1, one-cell-stage Tg(crx:gap-CFP) embryos were injected with 1 nl of hsp70:Stathmin1-mKate2 at 36 ng/uL ^45^, Tol2 transposase RNA at 80 ng/uL and GFP-ras RNA at 50ng/uL in ddH_2_O supplemented with 0.05% phenol red (Sigma-Aldrich). F_0_ embryos displaying high levels of GFP expression at 24 hpf were raised and germline carriers were identified by outcross with Tg(crx:gap-CFP) fish.

### DNA cloning and constructs used

The ath5:MRLC-mKate2 and ath5:GFP-UtrophinCH used to label myosin and actin, respectively, in PRs were assembled using Gateway cloning (Thermo Fisher Scientific) based on the Tol2 kit ^46^. The human MRLC2 gene from pCS2^+^-MRLC2-GFP ^47^ was used for creation of hsp70:MRLC-mKate2 construct which was then used as template for construction of MRLC2-mKate2 middle entry clone (pME) by PCR using Phusion polymerase (New England Biolabs) and primers with ATT recombination site (shown in lower case): Forward 5’-ggggacaagtttgtacaaaaaagcaggctggCTTCGCTGTCGTTTGTGGTCTCG-3’ and reverse 5’-ggggaccactttgtacaagaaagctgggtcTCATCTGTGCCCCAGTTTGC-3’. The pCS2^+^ vector containing GFP tagged calponin homology domain of human utrophin (GFP-UtrophinCH) ^48^ was used as template for generation of pME GFP- UtrophinCH by PCR using Phusion polymerase (New England Biolabs) and primers with ATT recombination site (shown in lower case): Forward 5’-ggggacaagtttgtacaaaaaagcaggctggATGGTGAGCAAGGGCGAGG and reverse ggggaccactttgtacaagaaagctgggtcTTAGTCTATGGTGACTTGCTGAG. The pME MRLC-mKate2 and pME GFP-UtrophinCH were combined with the 5′ entry clone containing the ath5 promoter ^46^ into the destination vector pTol2^+^pA R4-R2 backbone ^49^ to create the ath5:MRLC2-mKate2 and ath5:GFP-UtrophinCH.

The ath5:GFP-CAAX and trβ2:tdTomato were used to label PRs in wild-type embryos ^11,22^.The ath5:GFP-CAAX and ath5:GFP-DCX were used to label PRs expressing heat-shock inducible human Stathmin1(hsp70:Stathmin1-mKate2) ^11,45^. The βactin:GFP-DCX was used to label microtubules ^11^. The ath5:H2B-RFP was used to label the nucleus of PRs ^11^. The hsp70:H2B-RFP was used to label progenitor cells in colcemid treated embryos ^32^. The caggs:Lyn-tdTomato used as a transfection reporter in electroporation experiments was kindly provided by Takashi Namba (MPI-CBG, Dresden, Germany), modified version from Namba et al., 2014. The hcrx:GFP, gift from A. Swaroop (National Eye Institute, Bethesda, USA), containing conserved region of *crx* promoter of human origin was used to label PRs in human retinal organoids ^15^.

### DNA and morpholino injections of zebrafish embryos

To sparsely label cells or express proteins of interest in the zebrafish retina, one-cell-stage embryos were injected with purified plasmid DNA diluted in ddH_2_O supplemented with 0.05% phenol red (Sigma-Aldrich). Concentrations of individual constructs ranged from 10 to 25 ng/μL, but did not exceed 45 ng/μL when pooled. Injection volumes ranged from 0.5 to 1 nL. Morpholino targeting ath5 (5’-TTCATGGCTCTTCAAAAAAGTCTCC-3’, Gene Tools) was injected at 2 ng per embryo of Tg(ath5:gap-RFP) line to inhibit RGC generation ^27^. The p53 morpholino, 5’-GCGCCATTGCTTTGCAAGAATTG-3’ (Gene Tools), was co-injected at 2-4 ng per embryo with ath5 morpholino and hsp70:Stathmin1-mKate2 to reduce toxicity.

### Electroporation of retinal organoids

Labeling of human iPSCs-derived retinal organoids for live-imaging was done via electroporation with a transfection reporter (caggs:Lyn-tdTomato) and a photoreceptor reporter construct (hcrx:GFP). DNA preparation and electroporation procedure were adapted from previously published protocol for electroporation of mouse retina explants ^51^. Plasmids were purified with Endo Free Maxi Prep and a total amount of 100 μg of DNA (30% caggs:Lyn-tdTomato and 70% hcrx:GFP) was concentrated by precipitation with ethanol/sodium acetate and resuspended in 100 μL of sterile HBSS. Organoids were washed 2 times in HBSS prior to transfer to the electroporation chamber (petri dish with stainless steel electrodes embedded in silicone chamber of interior dimensions 0.4 × 0.5 × 0.8 cm—H × W × L). They were positioned close to the walls of the chamber to minimize deflation/compression during exchange of HBSS for 100 uL of 1μg/μl DNA mix. Next, organoids were positioned in the middle of the chamber with their largest surface facing the negative electrode, which was then electroporated. Using a square wave electroporation system (ECM 830 BTX), organoids were submitted to 5 square pulses of 25V for 50 ms with 950 ms intervals. Organoids were immediately washed 2 times in HBSS and transferred back to the appropriate cell culture medium.

### *In vivo* labelling of S-phase cells by EdU

To label S-phase cells, zebrafish embryos were incubated at 4°C for 1 h in E3 supplemented with 500 μm of EdU (ClickiT-Alexa 488 fluorophore kit, Invitrogen) while human retinal organoids were incubated at 37°C for 1h in organoid medium supplemented with 10 μm of EdU. Embryos and organoids were immediately fixed overnight in 4% PFA and, after antibody staining, incorporated EdU was detected according to manufacturer’s protocol.

### Drug treatments

Colcemid (Enzo Life Sciences), Rockout (Santa Cruz Biotechnology) and Blebbistatin (Enzo Life Sciences) were dissolved in DMSO and diluted to working concentrations in E3 medium. Same concentration DMSO was used in controls. For fixed imaging experiments, 30 hpf embryos were dechorionated and incubated at 32°C in multi-well plates with colcemid at 100 μM for 8 hours. For live imaging experiments, embryos embedded on agarose in compartmentalized 35-mm glass bottom Petri dishes (Greiner Bio-One) were incubated at 32°C with 25 and 100 μM of colcemid at 36 and 42 hpf, respectively; 50 μM of Rockout or 25 μM of Blebbistatin at 42 hpf until the end of imaging.

### Immunofluorescence

#### Whole-mount staining of retinal organoids

For validation of the specificity of the hcrx:GFP reporter ^15^, organoids co-electroporated with hcrx:GFP and caggs:Lyn-tdTomato at day 27 of culture were fixed overnight at day 43 in 4% PFA in PBS. Organoids were permeabilized with 0.5% Triton X-100 in PBS (PBS-T 0.5%) for 1 hour and blocked with 10% NDS in PBS-T 0.5% for 3 hours. Organoids were incubated for 2 days with primary antibodies (Anti-GFP at 1:100, A11120, Thermo Fisher Scientific; Anti-Otx2 at 1:100, HPA000633, Sigma-Aldrich) diluted in 1% NDS in PBS-T 0.2%. Next, organoids were washed 3 times for 30 min with PBS-T 0.2%, and incubated with appropriate fluorescently labelled secondary antibodies (Thermo Fisher Scientific) at 1:500 and RFP-Booster (rba594; Chromotek) at 1:200 for 2 days diluted in 1% NDS in PBS-T 0.2%. Organoids were washed 4 times for 20 min with PBS and stored in PBS at 4°C until imaging.

#### Whole-mount staining of zebrafish embryos

Zebrafish were dechorionated manually and fixed overnight in 4% PFA in PBS at 4°C. Embryos were washed 5 times for 15 min with PBS-T 0.8%. For permeabilization, embryos were incubated with 1x Trypsin-EDTA in PBS on ice for different periods of time depending on the developmental stage (10 min for 36 hpf, 12 min for 42 hpf and 15 min for 56 hpf). The permeabilization solution was replaced with PBS-T 0.8% and embryos were kept additional 30 min on ice before rinsing 2 times with PBS-T 0.8%. Blocking was performed with 10% NDS in PBS-T 0.8% for 3 h at room temperature. Embryos were incubated with primary antibodies diluted in 1% NDS in PBS-T 0.8% for 3 days at 4°C. Acetylated α-tubulin antibody (T6793; Sigma-Aldrich) 1:250, GFP antibodies (50430-2-AP, Proteintech; MAB3580, Millipore; Antibody Facility MPI-CBG) 1:250 and Histone H3 (phospho S28, ab10543, Abcam) 1:500. After 5 washes of 30 min with PBS-T 0.8%, embryos were incubated with appropriate secondary antibodies and DAPI in 1% NDS in PBS-T 0.8% for 3 days at 4 C. The RFP-Booster (rba594; Chromotek) was used at 1:200 and Rhodamine Phalloidin (R415, Thermo Fisher Scientific) was used at 1:50. Embryos were washed 4 times for 15 min with PBS-T 0.8% before storage in PBS at 4°C until imaging.

#### Tissue section staining: zebrafish, human fetal retinas and organoids

For immunostaining of sections of human fetal eyes, eyes were fixed in 4% PFA in PBS followed by cryopreservation in 15% and 30% sucrose in PBS. Zebrafish embryos and retinal human organoids were fixed in 4% PFA in PBS overnight at 4°C and cryopreserved in 30% sucrose in PBS overnight. Samples were embedded in OCT compound and transversal sections (10-15 μm) were collected on SuperFrost Plus slides (Thermo Fisher Scientific) and stored in −20°C. For immunofluorescence, sections were washed with PBS-T 0,2% for 5 min and submitted to antigenic retrieval when appropriate by boiling them in Citrate buffer 10mM pH 6.0 and letting it cool down at RT for at least 25 min. Sections were washed twice for 5 min with PBS-T 0,2%, permeabilize for 15 min with PBS-T 0,5% and washed for 5 min with PBS-T 0,2% prior to 45 min blocking with 10% NDS in PBS-T 0,2%. For human primary material, washes prior to the incubation with primary antibodies were performed with PBS without triton. Primary antibodies were incubated overnight or for 2 days at 4°C diluted in 1% NDS serum + 0,2% triton in PBS. CRX antibody (HPA036762; Sigma-Aldrich) 1:100, GFP antibody (Antibody Facility, MPI-CBG) 1:100, HuC/HuD antibody (A21271; Invitrogen) 1:250, OTX2 antibody (HPA000633; Sigma-Aldrich) 1:100, t Phospho-Myosin Light Chain 2 (Ser19) antibody (3671; Cell Signalling) 1:50 and Recoverin antibody (AB5585; Millipore) 1:100. Sections were washed three times 5 min with PBS-T 0,2% and incubated with appropriate fluorescently labelled secondary antibodies (Thermo Fisher Scientific) at 1:500 and DAPI for 2 hours at room temperature diluted in 1% NDS in PBS-T 0,2%. Sections were washed three times 5 min with PBS-T 0,2% and rinsed with PBS to remove Triton. Mounted slides with VectaShield (Vector Labs) were stored at −20°C.

### Image acquisition

#### Light-sheet imaging of retinal organoids

Electroporated retinal organoids were imaged using Lightsheet Z.1 (Carl Zeiss Microscopy) operated via ZEN 2015 (black edition) software. A syringe filter (0.22 μm, SLGV033RS, Millipore) was placed between the CO_2_ module and the chamber to minimize contamination. Electroporated organoids were pre-screened for expression of the transfection reporter using an inverted widefield fluorescence microscope (Leica DM IL LED) equipped with a 10x/0.25 NPLAN objective (Leica). Mounting procedure was adapted from previously published protocol for mounting of zebrafish embryos ^52^. In brief, molten 1% low melting point agarose (dissolved in PBS) was diluted to 0.6% with organoid medium (DMEM without phenol red/F12 3:1 supplemented with 2% B-27, 5% FBS, 1x GlutaMax, 1x NEAA, 0.1% Lipid concentrate, 1% Penicillin-Streptomycin and 100 μM Taurine). Up to three organoids were transferred to the agarose-medium mix and mounted by suction into a glass capillary (~1 mm inner diameter, Brand, #701904) by pulling an inserted Teflon tip plunger (Brand, #701932). The sample chamber was filled with organoid medium and maintained at 37°C and 5% of CO_2_. Single-view of each organoid spanning the electroporated region (~440 Z slices with step size of 1 μm) was acquired every 15 min for 18—35 hours using single sided illumination by 10X/0.2 objective (Carl Zeiss Microscopy). Detection was performed with Plan-Apochromat 20X/1.0 W objective (Carl Zeiss Microscopy) and two Edge 5.5 sCMOS cameras (PCO). For imaging of whole-mount stained organoids, they were embedded in 1% low melting point agarose (dissolved in PBS), the sample chamber was filled with PBS and imaging was performed at room temperature.

#### Light-sheet imaging of zebrafish embryos

Live imaging of wild type photoreceptors, cytoskeleton dynamics and genetic perturbations used Lightsheet Z.1 (Carl Zeiss Microscopy) operated via ZEN 2015 (black edition) software. Embryos were mounted in 0.6% low melting point agarose (prepared in E3) supplemented with 240 μg/mL of MS-222 (Sigma-Aldrich) as previously described ^52^. The sample chamber was filled with E3 medium supplemented with 120 μg/mL of MS-222 and 30.4 μg/mL of *N*-phenylthiourea and maintained at 28.5 C. Single-view spanning the entire volume of each eye (80-120 Z slices with step size of 1 μm) was acquired every 5 min for ~ 16 hours using dual sided illumination by 10X/0.2 objectives (Carl Zeiss Microscopy). Detection was performed with Plan-Apochromat 20X/1.0 W objective (Carl Zeiss Microscopy) and two Edge 5.5 sCMOS cameras (PCO).

#### Spinning disk imaging of zebrafish embryos

Live imaging of chemical (Colcemid, Blebbistatin and Rockout) and genetic (Tg(hsp70:Stathmin1-mKate2)) perturbations of cytoskeleton elements were performed using an Andor spinning disk confocal microscope composed of Andor IX 83 stand and a CSU-W1 scan head (Yokogawa) with Borealis upgrade. The microscope was operated via the Andor iQ software version 3.6. Embryos were embedded in 0.8% of low melting point agarose in E3 medium supplemented with 314 μg/mL of MS-222 and 0.1 M Hepes (pH 7.25) in compartmentalized 35-mm glass bottom Petri dish (Greiner Bio-One). The dish was filled with E3 supplemented with 120 μg/mL of MS-222 and 30.4 μg/mL of *N*-phenylthiourea and incubated at 32°C during imaging. Z-stacks of 60 μm with step size of 1 μm were acquired at 15 min intervals for 16 hours using Olympus UPLSAPO objective 60x 1.3 SIL and Andor iXon 888 Ultra with Fringe suppression.

#### Confocal scans of whole-mount stained zebrafish embryos

Whole-mount stained zebrafish embryos were imaged using Zeiss LSM 880 inverted point-scanning confocal microscope (Carl Zeiss Microscopy) operated via ZEN 2011 (black edition) software (Zeiss). Embryos were embedded in 1% of low melting point agarose (prepared in E3) in glass-bottom dishes (MatTek Corporation). The dish was filled with E3 and imaging was performed at room temperature using the 40x/1.2 C-Apochromat water immersion objective (Zeiss).

#### Confocal scans of tissue sections

Tissues sections mounted on microscope slides were imaged using Zeiss LSM 880 upright point-scanning confocal microscope (Carl Zeiss Microscopy) using 25x/0.8 LD LCI Plan-Apochromat and 63×/1.3 LCI Plan-Neofluar *multi-immersion* objectives (Zeiss). Microscope was operated via ZEN 2011 (black edition) software (Zeiss).

#### Image processing and analysis

Image data were processed in ZEN Black and/or Fiji ^53^. Spatial drift of live imaging data was corrected using Manual drift correction plugin from Fiji (by Benoit Lombardot, Scientific computing facility, MPI-CBG).

#### Deep learning-based image restoration

Tg(crx:gap-CFP) zebrafish line was imaged with minimal laser power and exposure using light-sheet (LSM) and spinning disk confocal (SDCM) microscopy. To improve signal-to-noise ratio, the time-lapse recordings were restored using the deep learning-based algorithm CARE ^20^. In brief, for each microscope setup (LSM and SDCM) a convolutional neural network was trained with pairs of low and high SNR stacks acquired at different imaging conditions (laser power and exposure time). For training and validation, we extracted 6885 patches of size (8 × 256 × 256) from 17 volumetric stacks for the LSM data, and 1280 patches of size (16, 128, 128) from 10 volumetric stacks for the SDCM data. For each volume acquisition conditions were intreleaved plane-by-plane to ensure proper spatial registration. A dedicated CARE network was trained for the LSM and SDCM restoration task, taking 15 hours 32 min and 10 hours and 51 min to converge, respectively. The python code for training data generation, CARE network training and prediction is available as Jupyter notebooks at [http://csbdeep.bioimagecomputing.com/doc/].

#### Cell tracking

For analysis of PR translocation in the zebrafish retina, wild-type and Stathmin1 overexpressing PRs labelled by mosaic injection of plasmids containing ath5-driven reporters (Fig. 3A and 3G; Extended Data Fig. 1C and 3F) and PRs in the ath5 knockdown condition labelled by Tg(ath5:gap-RFP) (Fig.3B) were identified based on reporter gene expression, morphology and final position at the apical surface of the retina. For analysis of PR translocation in the retinal human organoids, PRs labeled by electroporation with crx-driven reporter plasmid were identified based on the expression of the reporter gene (Fig. 2N and Extended Data Fig. 2D). Full trajectories from birth to final positioning and partial trajectories were obtained by tracing the center of the cell body in 2D maximum projected substacks of the raw data using ImageJ plugin MTrackJ ^54^.

#### Analysis of kinetics of migration

Mean square displacements (MSDs) and directionality ratios were calculated from the first 80 min of basal and apical migration of PRs in Excel (Microsoft) using the open-source computer program DiPer ^55^. To estimate directional persistence, the α-value was obtained from the slope of the log-log plots of MSDs and Time interval.

#### Analysis of the spatial distribution of PRs

The position of PRs cells with respect to the apical surface of developing human retinal organoids was measured manually using Fiji (Fig. 2L). The distance between the center of the cell body and the apical attachment was calculated in Excel.

#### Quantitative analysis of myosin enrichments

Myosin enrichments basal and apical to the nucleus of apically migrating PRs were measured in 2D sum projected substacks of the raw data. The 2D sum projections were registered to the center of the cell of interest using Fiji plugin for manual registration developed by the MPI-CBG Scientific Computing Facility. Value cutoff for thresholding was determined after median filtering (Median filter size: 1 pixel) by measuring the mean pixel intensity at the center of the cell (2×2 μm square) during the apical migration. The mean of the mean intensity of all timepoints plus two times the mean standard deviation was used as threshold limit. Values below the limit were set to NaN (Not a Number). The fraction of the positive area within the regions of interest (4×2 μm, Width × Height) basal and apical to the nucleus was measured for every timepoint of apical migration. The position of the square ROIs was adjusted manually according to the position of the cell, whereas the size of the ROIs remained constant.

#### Analysis of apical surface occupancy

To estimate the fraction of the apical surface area occupied by PRs at 42 hpf, the number of apical PRs and the total number of PRs were counted manually in subvolumes of the 42 hpf retina (N = 3 embryos). The tissue surface area in each subvolume was measured using a custom Fiji plugin (Volume Manager, by Robert Haase, Scientific computing facility, MPI-CBG) and visualized in 3D using Fiji plugin ClearVolume (Fig. 5B) ^56^. The cross-sectional area of apical PRs was measured at the center of the cell in the XZ orthogonal view (n= 11 cells, N = 4 embryos; Fig. 5C). To calculate the fraction of occupancy by apical PRs, the average cross-sectional area was multiplied by the number of apical PRs and subsequently divided by the tissue surface area. To calculate a theoretical occupancy in the case of no migration, the total number of PRs was used.

#### Statistical analysis

Statistical tests were performed using GraphPad Prism 6, except Dunn’s Kruskal-Wallis Multiple Comparisons with Holm-Bonferroni correction, which was conducted in python using scipy.stats v1.3.1 and v0.6.2 scikit-posthocs packages. Two-tailed tests were used and 95% confidence intervals were considered. See figure legends for more information on the P values and sample sizes.

#### Data availability

All data is available in the manuscript.

## Acknowledgments

We thank the Norden Lab for fruitful project discussions. We are grateful to E. Barriga, M. Cayouette, A. Grapin-Botton, W. Harris and C. Modes for helpful comments on the manuscript. We thank S. Kaufmann, H. Hollak, the Computer Department, and the Light Microscopy, Scientific Computing and Fish facilities of the Max Planck Institute of Molecular Cell Biology and Genetics for experimental support. We thank the Quantitative Biology unit at the Instituto Gulbenkian de Ciência for support with statistical analyses. We are grateful to T. Namba and A. Swaroop for sharing DNA constructs and thank M. Heide for the support with the human organoid culture. EN and JI are associated with the IMPRS-CellDevoSys PhD program and EN is also a member of the IBB-Integrative Biology and Biomedicine PhD program. This work was supported by MPI-CBG, the FCG-IGC, the German Research Foundation (NO 1069/5-1) and by an ERC consolidator grant (H2020 ERC-2018-CoG-81904).

## Author Contributions

MRM: Conceptualization, Data curation, Formal Analysis, Investigation, Methodology, Validation, Visualization, Writing – original draft, Supervision. JK: Formal Analysis, Investigation, Methodology, Validation. EN: Formal Analysis, Investigation, Methodology, Validation. MW: Software, Investigation, Methodology. IJ: Investigation, Methodology. EM: Resources, Supervision, Software. CN: Conceptualization, Funding acquisition, Project administration, Resources, Writing – original draft, Supervision.

## Competing Interests

The authors declare no competing financial interests.

**Supplementary Information** is available for this paper.

---

**Extended Data Fig. 1.**
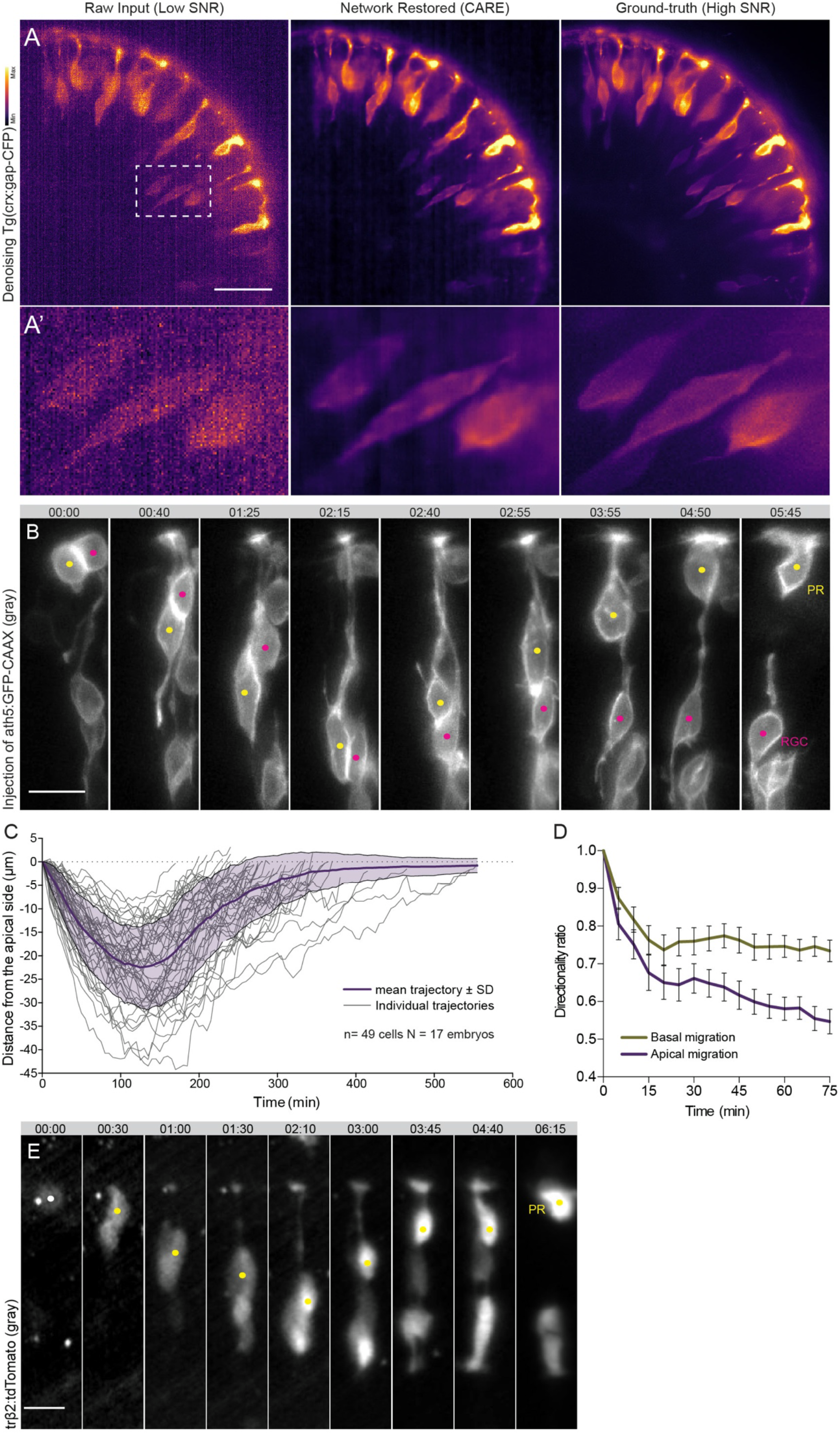
(A) Example of denoising of the crx:CFP signal (gray) using the deep learning-based algorithm CARE. Dashed white box indicates areas shown in A’. (B) Bidirectional migration of photoreceptor (PR). ath5:GFP-CAAX (gray) labels early neurogenic progenitors; Time is displayed in hours:minutes. Yellow dots mark PR; Magenta dots mark RGC. (C) Trajectories of PR bidirectional migration from birth to final positioning relative to the apical surface (0 μm) shown as individual trajectories (n= 49 cells, N= 17 embryos) with superimposed mean trajectory ± SD. (D) Directionality ratio of PR’s basal and apical migration shown as mean of all the tracks ± SEM (n= 33 cells; N= 15 fish). (E) Bidirectional migration of photoreceptor (PR). trβ2:tdTomato (gray) labels L-cone PRs; Time is displayed in hours:minutes. White dot marks apical birth; Yellow dots mark PRs. Scale bars: 10 μm (B, E), 25 μm (A).

**Extended Data Fig. 2.**
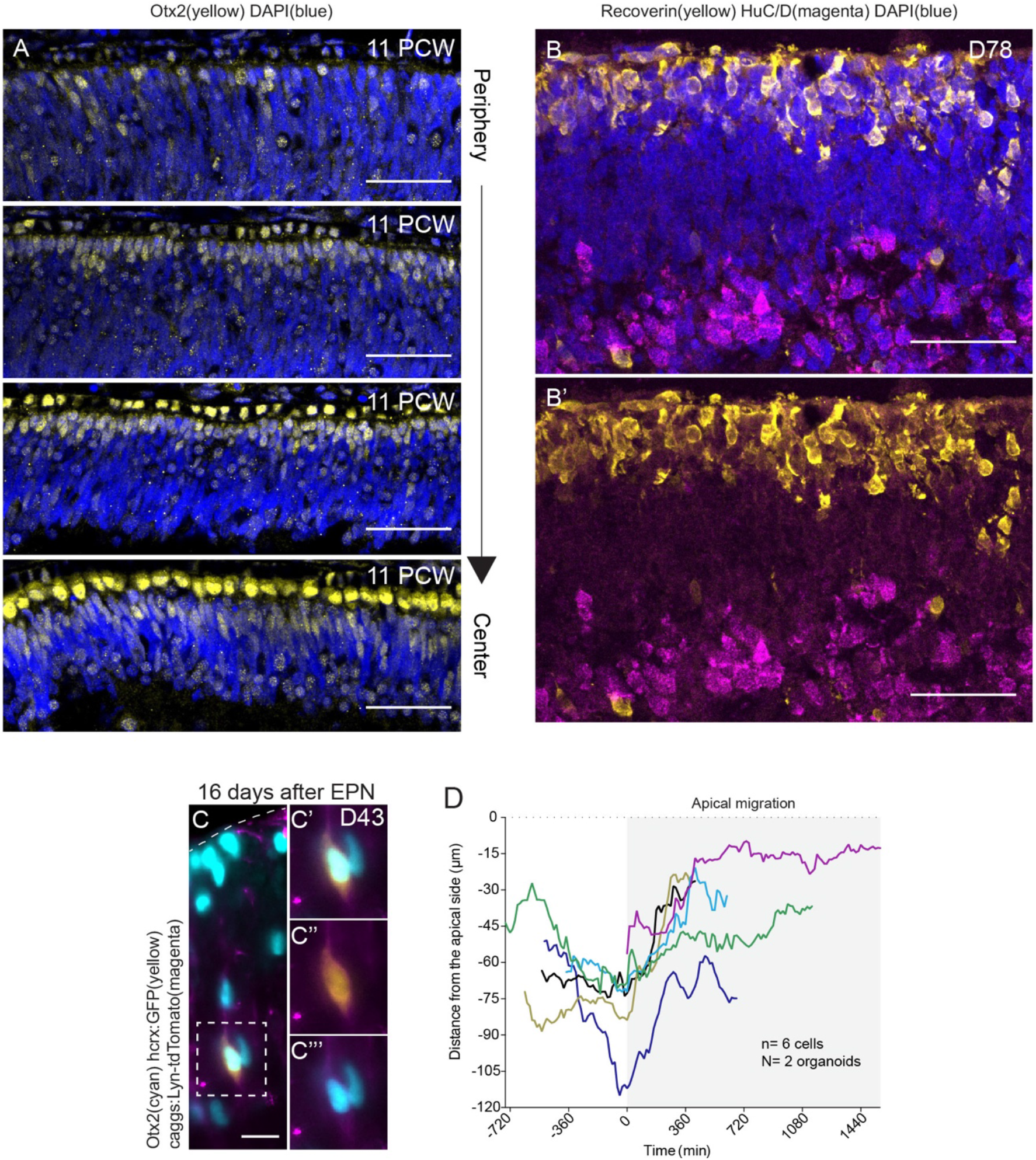
(A) Spatial distribution of Otx2+ PRs along the apical-basal axis of the neuroepithelium of 11 PCW human retina from peripheral to central regions. Staining: Otx2 (yellow) labels Otx2+ PRs; DAPI (blue) labels nuclei. (B) Spatial distribution of Recoverin+ cells along the apical-basal axis of the neuroepithelium of human retinal organoids at an early stage of lamination (D78). Staining: Recoverin (yellow) labels Recoverin+ PRs; HuC/D (magenta) labels RGC and amacrine cells; DAPI (blue) labels nuclei. (C) Identity of cells expressing GFP under the control of the human *crx* promoter at 16 days after electroporation (12 Otx2+ of 13 GFP+ cells analyzed, N= 5 organoids). Staining: endogenous Otx2 (cyan); hcrx:GFP (yellow); caggs:Lyn-tdTomato (magenta). (D) Partial trajectories of migration of PRs positive for hcrx-driven GFP relative to the apical surface (0 μm) and the start of apical migration (0 min). Scale bars: 20 μm (C), 50 μm (A-B).

**Extended Data Fig. 3.**
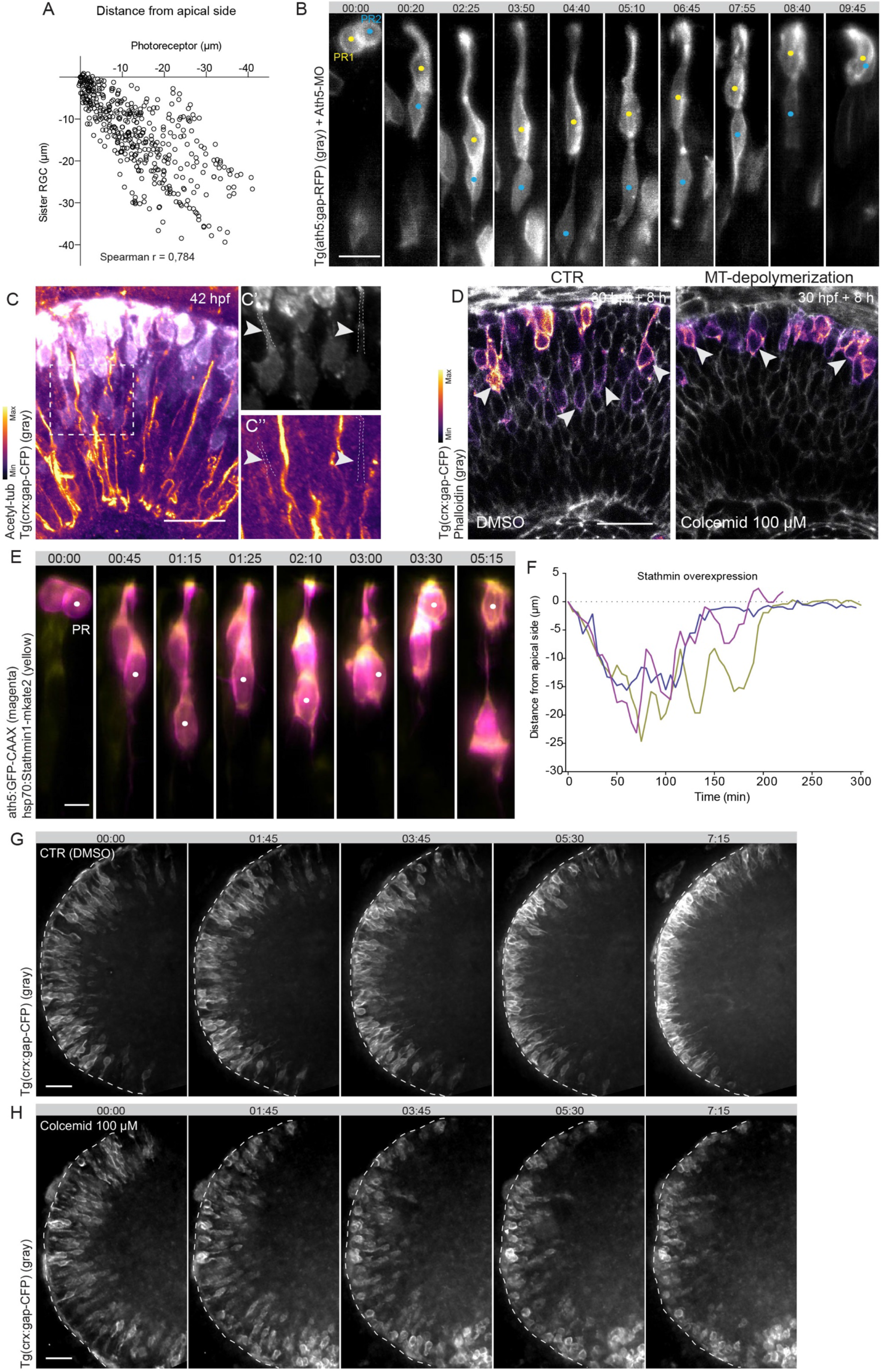
(A) Correlation between PR and sister RGC positions relative to the apical surface (0 μm). Each dot represents an individual timepoint from 20 pairs of PR-RGC cells (450 timepoints, N= 7 embryos). (B) Bidirectional migration of sister PRs upon Ath5 morpholino mediated knockdown. Tg(ath5:gap-RFP) (gray) labels early neurogenic progenitors. Time is displayed in hours:minutes. Yellow and blue dots mark sister PRs. (C) Distribution of stable MTs in apically migrating PRs at 42 hpf. Staining: Tg(crx:gap-CFP) (gray) labels PRs; Acetylated tubulin labels stable MTs, lookup table shows min and max signal values. Dashed white box indicates areas shown in C’ and C’’. Arrowheads mark basal process of PRs. (D) Effect of colcemid-induced MT depolymerization on PR basal migration. Embryos were treated with 100 μM of colcemid (DMSO for control) from 30 hpf for 8 h. Staining: Tg(crx:gap-CFP) labels PRs, lookup table shows min and max signal values; phalloidin (gray) labels F-actin. Arrowheads mark PRs. (E-F) Effect of Stathmin1 overexpression on PR migration. (E) ath5:GFP-CAAX (magenta) labels early neurogenic progenitors; hsp70:Stathmin1-mKate2 (yellow). White dots mark PR. (F) Trajectories of PRs upon Stathmin1 overexpression relative to the apical surface (0 μm). (G-H) Effect of colcemid-induced MT depolymerization on PR apical migration. Time-series of retinas treated with colcemid (100 μM, H) or DMSO (control, G). Tg(crx:gap-CFP) (gray) labels PRs. Time is displayed in hours:minutes. Scale bars: 5 μm (E), 10 μm (B), 20 μm (C-D, G-H).

**Extended Data Fig. 4.**
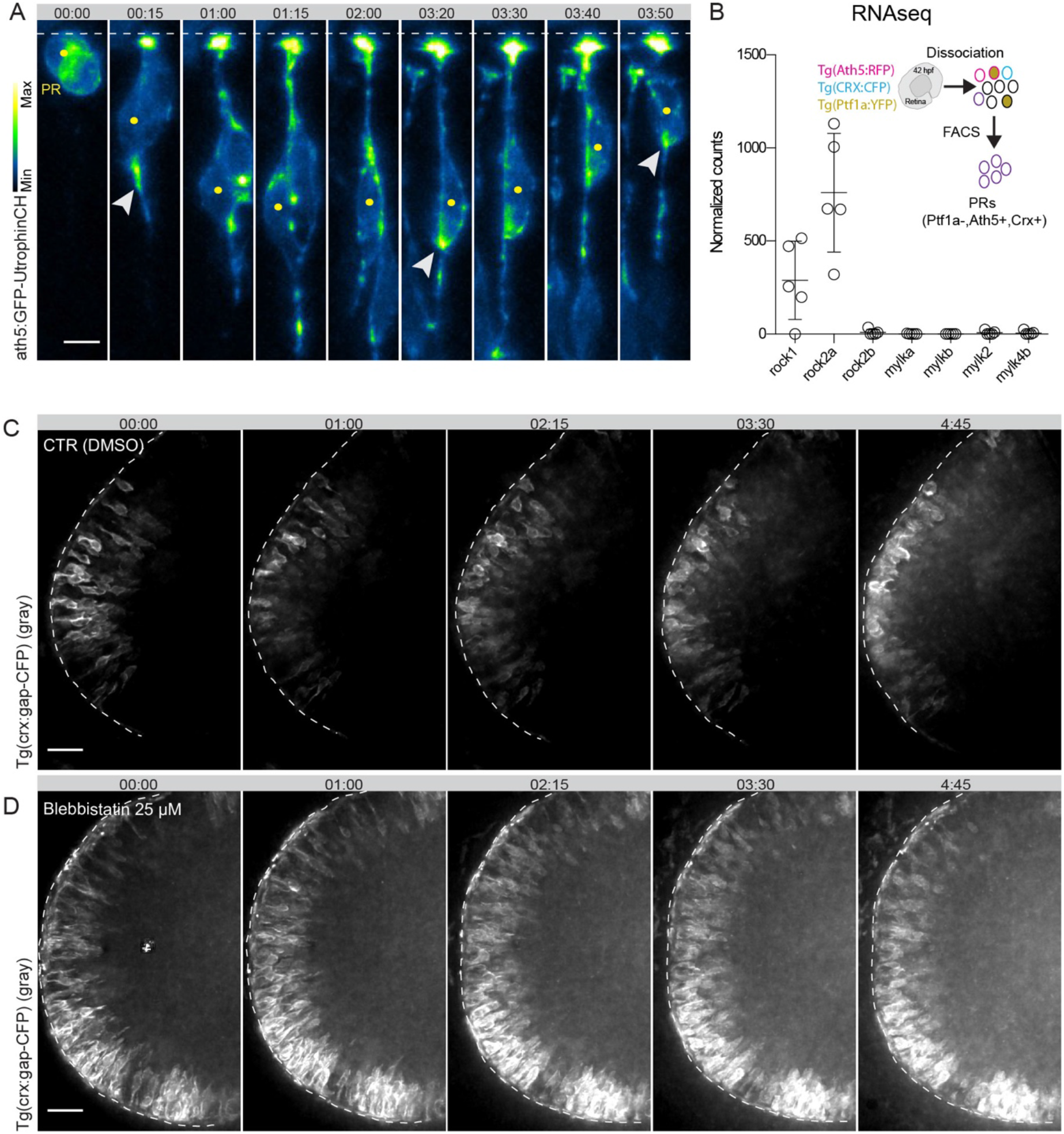
(A) Actin distribution during PR translocation. ath5:Utrophin-GFP labels actin in early neurogenic progenitors, lookup table shows min and max signal values. Time is displayed in hours:minutes. Yellow dots mark PR; arrowheads mark actin enrichment. (B) Expression levels of *rock* and *mylk* genes in PRs at 42 hpf shown as mean ± SD (N=5 embryos). (C-D) Time-series of retinas treated with Blebbistatin (25 μM, D) or DMSO (control, C). Tg(crx:gap-CFP) (gray) labels PRs. Time is displayed in hours:minutes. Scale bars: 5 μm (A), 20 μm (C-D).

**Extended Data Fig. 5.**
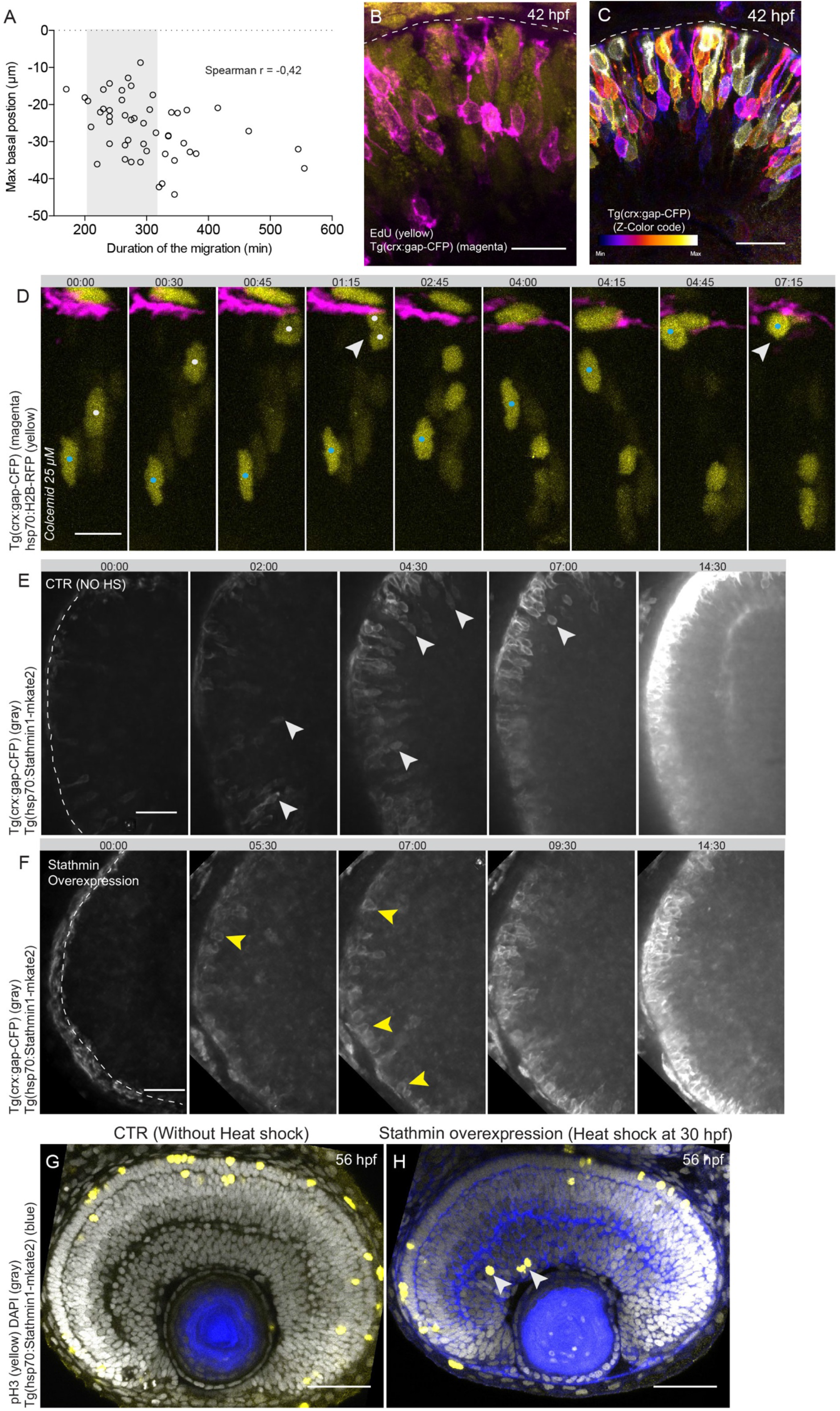
(A) Correlation between max basal position and duration of photoreceptor (PR) migration. Each dot represents individual cells (n= 49 cells, N= 17 embryos). (B) Proliferative status of the retina during PR migration at 42 hpf. Staining: Tg(crx:gap-CFP) (magenta) labels PRs; EdU (yellow) labels cell in S-phase. (C) Abundance of basal PRs at 42 hpf. Tg(crx:gap-CFP) labels PRs, color represents z-depth, lookup table shows min and max z depths. (D) Progenitor movements and apical mitosis upon colcemid treatment in an area in which PRs did not yet emerge. Embryos were treated with 25 μM of colcemid. Tg(crx:gap-CFP) in magenta labels PRs; hsp70:H2B-RFP in yellow labels apically migrating nuclei. Time is displayed as hours:minutes. Blue and white dots mark progenitor cells; arrowheads mark apical mitosis. (E-F) Time-series of photoreceptor lamination upon Stathmin-overexpression induced blockage of PR migration (F) and control retina without heat shock (E). Heat shock induction of Stathmin expression at 36 hpf. Tg(crx:gap-CFP) (gray) labels PRs. Time is displayed in hours:minutes. White arrowheads mark basal PRs; Yellow arrowheads mark apical PRs. (G-H) Effect of Stathmin-induced blockage of PR migration on progenitor divisions at 56 hpf. Heat shock induction of Stathmin overexpression at 30 hpf. Staining: Tg(hsp70:Stathmin1-mkate2) (blue) labels Stathmin, pH3 (yellow) labels mitotic cells; DAPI (gray) labels nuclei. Arrowheads mark ectopic mitosis. Scale bars: 10 μm (D), 15 μm (B), 20 μm (C), 25 μm (E-F), 50 μm (G-H).

**Extended Data Fig. 6.**
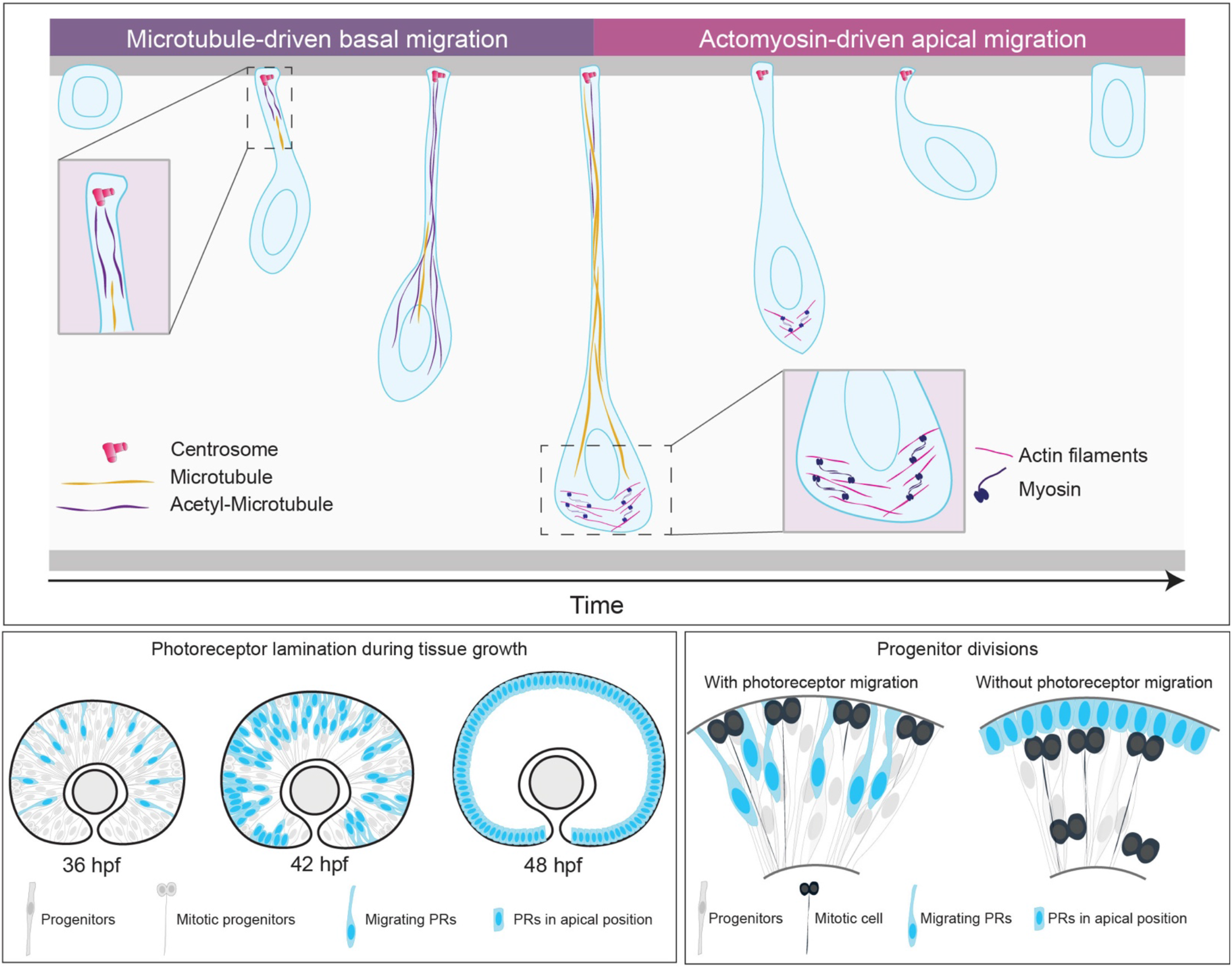
Schematic summary of mechanisms driving photoreceptor migration and its relevance for the coordination of tissue growth and organization. (Upper panel) Photoreceptors (PR) emerge at the apical surface of the retinal neuroepithelium and undergo bidirectional somal translocation that entails fast and directed movement towards the basal lamina followed by a basal pause and saltatory movements towards the apical surface. Basal and apical migration display different kinetics due to the involvement of distinct cytoskeleton machineries. While basal migration depends on stable microtubules, they are dispensable for apical translocation, which depends on actomyosin contractions at the cell rear. (Lower left panel) PR bidirectional migration is concurrent with cell proliferation and tissue growth. (Lower right panel) Bidirectional migration of PRs delays lamination helping to coordinate tissue growth and organization by securing space at the apical surface for incoming divisions of progenitor cells. This bidirectional mode of somal translocation is crucial to maintain tissue integrity and function, as blockage of PR movements leads to overcrowding of the mitotic zone and subsequent progenitor delamination.

## Supplemental video legends

**Supplementary Video 1** Bidirectional migration of photoreceptors in the zebrafish retina, related to Fig. 1. (Part 1) Tg(ath5:gap-RFP) (gray) labels early neurogenic progenitors; Tg(crx:gap-CFP) labels PRs, lookup table shows min and max signal values. (Part 2) trβ2:tdTomato (gray) labels L-cone PRs. Time is displayed in hours:minutes. Yellow dot marks PRs.

**Supplementary Video 2** Photoreceptor layer formation in the zebrafish retina, related to Fig.

1. Tg(ath5:gap-RFP) (gray) labels early neurogenic progenitors; Tg(crx:gap-CFP) labels PRs, lookup table shows min and max signal values. Time is displayed in hours:minutes.

**Supplementary Video 3** Bidirectional migration of photoreceptors in human retinal organoids, related to Fig. 2. hcrx:GFP (yellow) labels PR; caggs:Lyn-tdTomato (magenta) labels transfected cells. Blue dots mark PRs. Time is displayed in hours:minutes.

**Supplementary Video 4** Effect of Stathmin1 overexpression on photoreceptor migration, related to Fig. 3. ath5:GFP-CAAX (magenta) labels early neurogenic progenitors; hsp70:Stathmin1-mKate2 (yellow). Blue dots mark PR. Part 1 shows complete stalling of PR translocation. Part 2 shows loss of directionality phenotype. Time is displayed in hours:minutes.

**Supplementary Video 5** Effect of tissue-wide interference with microtubules or myosin on photoreceptor migration, related to Fig. 3, 4 and 5. (Part 1) Effect of Stathmin overexpression resulting in MT depolymerization on PR basal migration. Heat shock induction of Stathmin expression at 36 hpf. Tg(crx:gap-CFP) (gray) labels PRs, Tg(hsp70:Stathmin1-mKate2) in blue. (Part 2) Effect of colcemid-induced MT depolymerization on PR apical migration. Time-series of retinas treated with colcemid (100 μM) or DMSO (control). Tg(crx:gap-CFP) (gray) labels PRs. (Part 3) Effect of myosin inhibitor Blebbistatin on PR apical migration. Time-series of retinas treated with Blebbistatin (25 μM) or DMSO (control). Tg(crx:gap-CFP) (gray) labels PRs. (Part 4) Effect of actomyosin activity inhibitor Rockout on PR apical migration. Time-series of retinas treated with Rockout (50μM) or DMSO (control). Tg(crx:gap-CFP) (gray) labels PRs. Time is displayed in hours:minutes.

**Supplementary Video 6** Myosin distribution during photoreceptor migration, related to Fig. 4. Myosin distribution during PR migration. ath5:GFP-CAAX (magenta) labels early neurogenic progenitors; ath5:MRLC2-mKate2 labels myosin, lookup table shows min and max signal values. Yellow dot marks PR; arrowheads mark myosin enrichments. Time is displayed in hours:minutes.

**Supplementary Video 7** Effect of colcemid-induced blockage of PR migration on progenitor divisions, related to Fig. 5. (Part 1) Subapical mitoses upon colcemid-induced blockage of basal PR migration. (Part 2) Progenitor movements and apical mitosis upon colcemid treatment in an area in which PRs did not yet emerge. Embryos were treated with 25 μM of colcemid. Tg(crx:gap-CFP) in magenta labels PRs; hsp70:H2B-RFP in yellow labels apically migrating progenitor nuclei. Blue and white dots mark progenitor cells. Time is displayed in hours:minutes.

